# The chemorepellent effect of metformin limits bacterial penetration into mammalian mucus by altering bacterial motility

**DOI:** 10.64898/2026.09.17.752276

**Authors:** Kevin Simpson, Axel Ranson, Marta Vazquez Gomez, Alex Le Guen, Aline Rifflet, Richard Wheeler, Héloïse Rytter, Clara Delaroque, Guy Gorochov, Benoit Chassaing, Tiphaine Le Roy, Ivo Gomperts Boneca, Éric Clément, Karine Clément

## Abstract

Metformin improves metabolic health and remodels the intestinal microbiota, yet how these microbial changes contribute to therapeutic benefits remains incompletely understood. We combined human metagenomics, quantitative analyses of bacterial behavior and ex vivo mucus models to investigate how metformin modulates host–microbiota interactions. Human metagenomics revealed metformin-associated enrichment of chemotaxis- and motility-related genes. At millimolar concentrations, metformin acted as a bacterial chemorepellent, an effect abolished by deletion of the chemotaxis regulator CheY and dependent on the chemoreceptor Tsr. Direct metformin exposure or fecal water from metformin-treated individuals reduced bacterial penetration into mucin, an effect reproduced in intestinal mucus from pigs fed a moderate-fat diet. Metformin treatment was further associated with reduced fecal flagellin activity and increased anti-flagellin IgA. Together, these findings identify bacterial chemotaxis as a target of metformin and reveal a mechanism by which metformin may limit bacterial access to the intestinal mucus barrier.

## Introduction

Obesity and type 2 diabetes (T2D) are associated with alterations of the gut microbiota that contribute to metabolic dysfunction and chronic low-grade inflammation ^1–3^. These diseases arise from complex interactions between genetic susceptibility and environmental factors, among which diet and lifestyle play central roles, with gut microbiota acting as both a mediator and an effector of host metabolism and inflammation ^4–8^.

Earlier studies described taxonomic signatures of dysbiosis, but accumulating evidence indicates that microbial functional capacity, including nutrient metabolism, bile acid transformation and immune modulation, to cite a few, may better explain metabolic phenotypes than community composition alone ^7,9,10^. An additional and largely overlooked dimension is the spatial organization of the gut microbiota.

Beyond the composition and functional capacities of resident microorganisms, their physical proximity to the intestinal epithelium shapes host–microbe interactions. In obesity and T2D, disruption of this spatial segregation allows bacteria to migrate closer to the epithelial surface, a phenomenon previously termed bacterial encroachment ^11–13^. The intestinal mucus barrier is the primary structure maintaining this spatial segregation between the microbiota and the host ^14–18^. Disruption of this barrier facilitates bacterial encroachment and host exposure to immunostimulatory products such as lipopolysaccharide and peptidoglycan, thereby promoting chronic low-grade inflammation and metabolic dysfunction ^13,19–21^. Preserving microbiota–host spatial segregation has therefore emerged as an important determinant of metabolic homeostasis.

Whether currently available therapies influence microbiota–host spatial organization remains to be explored. Metformin, the first-line treatment for T2D, accumulates within the intestinal lumen at concentrations substantially exceeding circulating levels and has emerged as one of the most reproducible pharmacological modulators of the gut microbiota, also suggesting that part of its therapeutic efficacy may be mediated through the intestinal ecosystem rather than the host alone ^22–27^. Across multiple cohorts, metformin reshapes microbiome community composition, notably through depletion of *Roseburia* spp., enrichment of *Akkermansia muciniphila,* reduces microbial diversity and enhances the relative abundance of Proteobacteria, a taxonomic shift traditionally associated with dysbiosis ^23–26,28–30^.

Beyond taxonomic composition, metformin remodels microbial functional pathways, including metabolite production and flagellar biosynthesis ^26,31–33^. Because flagellin governs both bacterial motility and TLR5-mediated anti-flagellin IgA responses, these changes could influence bacterial localization at the mucosal surface and host–microbiota interactions ^34–37^. Yet, whether metformin directly alters bacterial behavior and access to the mucus barrier has not been experimentally demonstrated. In parallel, metformin has been reported to enhance mucus barrier function, further suggesting coordinated effects on both the microbiota and the host interface ^38^.

Together, these observations reveal an apparent paradox. Although metformin improves metabolic health, it also induces changes in microbial phenotype and function, including reduced diversity, increased Proteobacteria abundance and enrichment of motility-related functions, that are traditionally associated with metabolic dysfunction and impaired barrier integrity. This discrepancy suggests that microbial composition alone may not capture the functional consequences of metformin treatment and raises the possibility that bacterial behavior and spatial organization are important determinants of microbiota–host interactions. How these apparently contradictory observations are reconciled remains unknown. Specifically, it remains to be determined whether metformin-induced changes in bacterial motility translate into altered bacterial behavior at the mucosal surface, thereby limiting bacterial penetration and host exposure to microbial products, or instead, promote bacterial encroachment despite favorable metabolic outcomes.

Here, we combined human metagenomics with quantitative analyses of bacterial behavior and *ex vivo* mucus and mucin models to investigate how metformin influences bacterial motility and spatial interactions with the intestinal mucus barrier. We specifically examined whether metformin-associated changes in microbial motility functions translate into altered bacterial behavior and whether direct exposure to metformin affects bacterial penetration into mucus. By integrating human microbiome data with mechanistic bacterial and mucus models, we sought to determine whether bacterial behavior represents an additional dimension of the microbiota response to metformin.

## Results

### Metformin treatment is associated with enrichment of bacterial chemotaxis and mobility functions

French participants of the MetaCardis cohort were stratified into five groups based on metabolic status and metformin treatment lean metabolically health subjects (Lean-MH), metabolically healthy participants with obesity (Ob-MH), metabolically unhealthy patients with obesity (Ob-MUH) and participants with obesity and type 2 diabetes (T2D) without metformin treatment (Ob-T2Dna) or participants with obesity and T2D with metformin treatment (Ob-T2Dmtf). Detailed clinical characteristics are provided in Table S1. As expected, obesity and T2D were associated with a graded deterioration of metabolic, hepatic and inflammatory profiles compared with lean healthy individuals, while metformin-treated subjects displayed a partially improved inflammatory profile.

We first examined the overall structure and composition of the gut microbiome, using α-diversity and β-diversity ^39^. Compared with the Lean-MH group, all groups with metabolic alterations exhibited lower α-diversity, albeit with a more pronounced reduction in bacterial diversity in the Ob-T2Dmtf group showing the lowest values for both richness and Shannon index (Fig 1A, Fig S1A, Table S3A). Analysis of β-diversity further revealed differences in community structure between groups, with the Lean-MH group and the Ob-T2Dmtf group showing the most distinct profiles, whereas groups with untreated metabolic alterations displayed more overlapping compositions (Fig 1B, Fig S1B, Table S3B). At the phylum level, the Ob-T2Dmtf group was characterized by higher relative abundances of Bacteroidota and Proteobacteria, notably *E. coli*, and lower relative abundances of Firmicutes compared with the other groups (Fig 1B, Fig S1C).

**Fig 1:**
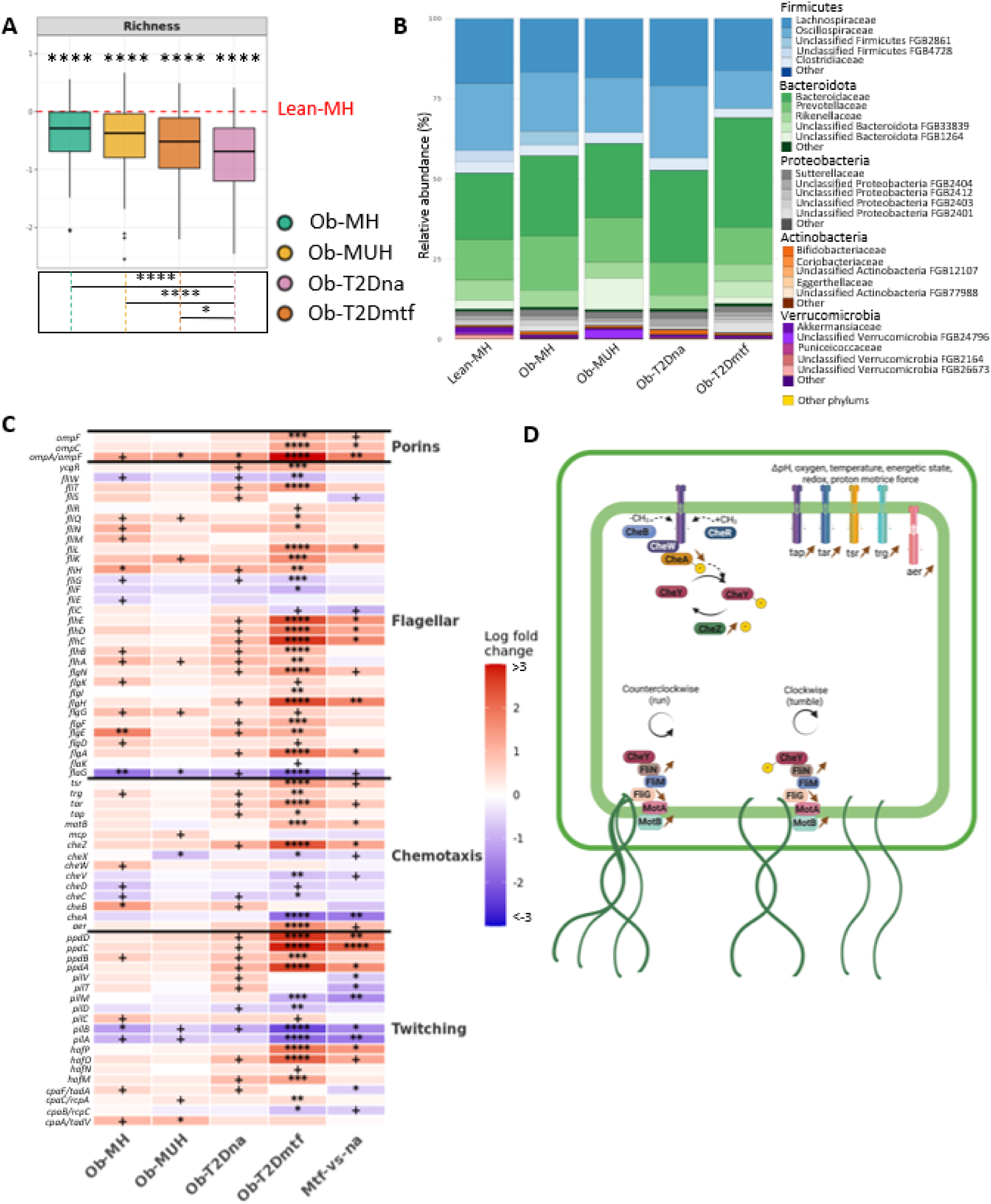
Alteration of bacterial composition and structure by metabolic diseases and metformin treatment. **A:** Log-transformed fold change in alpha diversity (richness) relative to the Lean-MH group. The dashed red line indicates the Lean-MH reference value (log fold change = 0). Pairwise comparisons were performed using Wilcoxon rank-sum tests, and p-values were adjusted for multiple testing using the Benjamini–Hochberg (BH) procedure. Adjusted False Discovery rate (FDR) values (***) above the plot correspond to comparisons of each clinical group to the Lean-MH group, whereas *** in the box below correspond to comparisons between diseased groups. **B:** Stacked bar plots showing microbial composition across groups. The plots represent the mean of the relative abundance of the five most represented families within the five most abundant phyla: Firmicutes (blue), Actinobacteria (orange), Bacteroidota (green), Proteobacteria (grey) and Verrucomicrobia (purple). Remaining families and phyla are grouped under “Other families” and “Other phyla” (yellow). **C:** Heatmap showing differences in the relative abundance of bacterial motility–related KEGG orthologs (KOs), grouped by motility category, identified using MaAsLin2, adjusted for age and gender. For the first four columns, groups are compared with the Lean-MH group; the last column shows the comparison between Ob-T2D participants treated or not treated with metformin. In blue: Decrease in gene abundance. In red: Increase in gene abundance, represented by the log of the fold change. **D:** Diagram showing how a Gram-negative bacterium recognizes its external environment through the MCP and chemotaxis pathways. The arrows indicate a significant difference in the abundance of the genes between the Ob-T2Dmtf group and the Lean-MH group. Lean-MH: lean metabolically healthy, Ob-MH: obesity and metabolically healthy, Ob-MUH: obesity and metabolically unhealthy, Ob-T2Dna: obesity and type 2 diabetes without metformin and Ob-T2Dmtf: obesity and type 2 diabetes treated by metformin. +: p ≤ 0.05, *: FDR ≤ 0.05, **: FDR ≤ 0.01, ***: FDR ≤ 0.001, ****: FDR ≤ 0.0001.

We next centered our observations on the examination of microbial functional potential, using KEGG Orthology (KO) to examine differences in gene enrichments between the clinical groups (Table S2). Interestingly, we identified significant differences between the abundance of multiple motility- and environmental sensing-related gene families, including genes involved in flagellar assembly (e.g., *fliC*, *flgM*), environmental sensing and methyl-accepting chemotaxis proteins (MCPs) (e.g., *tsr*, *tar*, *trg*, *tap*, *aer*), chemotaxis signaling components (e.g., *cheA*, *cheZ*), and pili-associated twitching motility (Fig 1C, Table S5).

Following normalization of sequencing depth across groups, the metformin-treated group exhibited a distinct distribution of gene abundances compared with the other groups (Fig 1C, Table S5). First, genes involved in type IV pilus assembly were significantly depleted in the Ob-T2Dmtf group (Fig 1C). Given the established role of type IV pili in twitching motility, bacterial attachment and early biofilm formation, this depletion may reflect a reduced representation of these functions within metformin-associated microbial communities ^40,41^. These changes may therefore indicate broader alterations in bacterial motility and surface interactions at the host–microbiota interface, consistent with previous observations reported in metformin-treated populations.

Moreover, metformin-treated individuals exhibited a higher abundance of genes encoding methyl-accepting chemotaxis proteins (MCPs), *aer* and *cheZ*, together with a lower abundance of *cheA* (FDR ≤ 0.0001). The proteins encoded by these genes are components of canonical bacterial chemotaxis pathway (sketched in Fig. 1D), which senses environmental gradients and controls run-and-tumble behavior and directional bacterial migration. This system has been extensively characterized in peritrichous *E. coli* ^42^.

At the level of the auto-phosphorylating histidine kinase CheA, the release of phosphoryl groups (P), in the presence of the response regulator CheY, produces CheY-P that binds to FliM, a component of the rotor protein assembly. This binding consequently switches the motor rotation, which induces a disassembling process of the flagellar bundle, hence changing the bacterium swimming direction ^43,44^. Conversely, CheZ dephosphorylates CheY-P, thereby reducing the motor rotation switch rate. These findings suggest an altered representation of chemotaxis-related genetic functions within metformin-associated microbial communities consistent with increased run persistence ^45–47^. Moreover, genes associated with the flagellar system were also generally enriched, with only a limited number of genes showing reduced abundance. Among them *fliC*, encoding flagellin protein, was significantly depleted (p ≤ 0.05) (Fig 1C). As flagellin is both the major structural component of the flagellum and the principal ligand recognized by TLR5, this pattern highlights a differential distribution of flagellar-related genetic functions in metformin-treated individuals. Finally, the increased abundance of OmpC- and OmpF-associated functions may also reflect enhanced genetic potential for solute exchange and outer membrane homeostasis in a metformin-exposed environment ^48,49^.

Overall, metformin treatment was associated with coordinated changes in the abundance of genes involved in chemotaxis, motility and membrane transport. However, metagenomic analyses capture genetic potential, rather than bacterial behavior, and therefore cannot determine whether these signatures translate into altered motility or host–microbiota interactions. We therefore used the motile *E. coli* strain AD62, a representative Proteobacterium enriched in metformin-treated individuals, to directly test the effects of metformin on bacterial movement.

### Metformin alters bacterial motility and directional behavior *in vitro*

To assess the functional relevance of the metagenomic signatures associated with metformin exposure, we quantified flagellin activity, bacterial motility parameters and directional responses to metformin *in vitro* using concentrations spanning the range measured in fecal water from study participants (Fig 2A–C, Fig S2). LC-MS/MS analysis detected metformin at millimolar concentrations in fecal samples, confirming substantial intestinal exposure to the drug (Fig S2).

**Fig 2:**
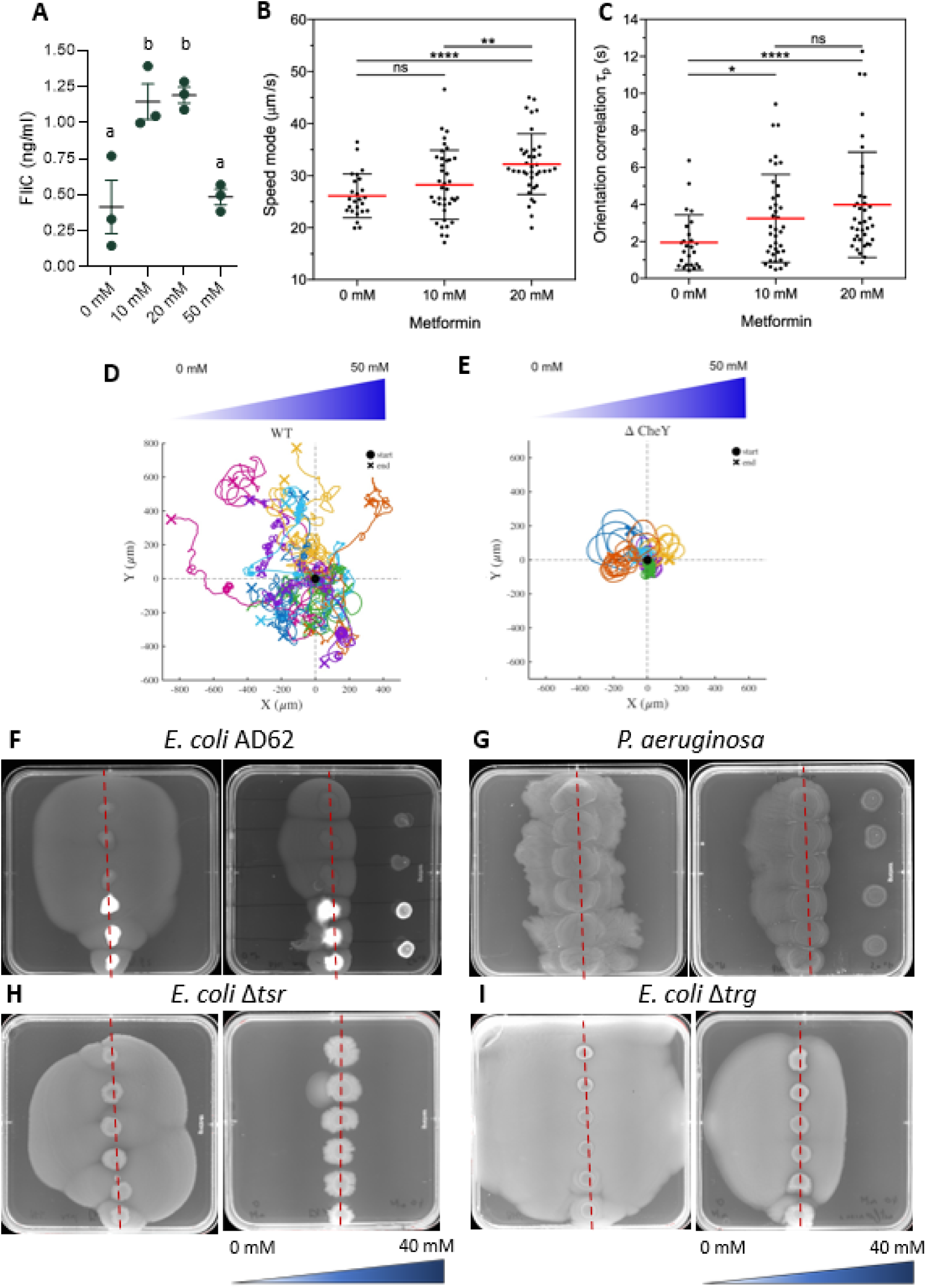
Metformin impacts bacterial motility in vitro. **A:** Flagellin bioactivity measured using a TLR5 reporter assay in supernatants from *E. coli* cultures exposed to increasing concentrations of metformin. Each biological replicate corresponds to an independent bacterial culture, performed in triplicate. For each biological replicate, metformin exposure was carried out in technical triplicate, and flagellin bioactivity was subsequently quantified using TLR5-HEK reporter cells in technical triplicate. Points represent biological replicates and bars indicate mean ± SEM. Statistical differences between metformin concentrations were assessed using one-way ANOVA followed by Tukey’s multiple-comparisons test. Groups sharing the same letter are not significantly different after p-value adjustment, whereas groups with different letters differ significantly (Tukey-adjusted p-value ≤ 0.05). **B-C**: Measurement of the swimming speed (B) and swimming orientation persistence time (C) of *E. coli* in a liquid culture with different concentrations of metformin. Each data point corresponds to a single bacterium. Statistical differences were calculated based on non-parametric, unpaired two-tailed Mann-Whitney test (α = 5%). ns (not significant): p > 0.05, *: p ≤ 0.05, **: p ≤ 0.01, ****: p ≤ 0.0001. **D-E**: Trajectories of WT (AD62) (D) or *cheY*-deleted (AD63) *(E) E. coli* in liquid medium with a metformin concentration gradient, from 0 to 50 mM. Each line represents the movement of a single bacterium, and the cross indicates its position at the end of the tracking period. The displacements along the horizontal direction were monitored for a time duration of *T* = 120 s. Then the tracks were shifted so that they began at the same starting position (0,0). **F-I**: Bacterial migration in 0.3% agar onto 1% agar without (left) or with (right) a metformin gradient ranging from 0 to 40 mM for WT *E. coli* AD62 (F), *P. aeruginosa* (G), *E. coli* Δ*tsr* (H), and *E. coli* Δ*trg* (I). The vertical red dotted lines indicate the location where the bacterial droplets were deposited in the center of the Petri dishes. Strains were also inoculated in the area of highest concentration of metformin to study the response at high concentrations for F-G.

Using a TLR5 reporter assay, we observed that metformin increased bioactive flagellin production by *E. coli* over a range of concentrations, up to 50 mM, above which bacterial growth became significantly impaired (Fig 2A, Fig S3A). Although this observation differs from the lower abundance of *fliC* detected in metagenomic analyses, these two measurements capture distinct biological levels, namely gene abundance within microbial communities versus flagellin activity by an individual bacterial strain under controlled experimental conditions.

We next tested whether the metformin-associated metagenomic signature may translate into change in bacterial behavior in a controlled *in vitro* experiment using a Lagrangian tracking system. This tool is suitable to monitor the exploration of a single bacterium in 3D ^50,51^ and we evaluated the bacterial velocity and trajectory using a dose-dependent exposure to metformin as well as the chemotactic response to a metformin gradient (Fig 2B-E).

First, we observed that metformin moderately increased *E. coli* motility in liquid, resulting in an approximately 1.2-fold increase in swimming speed at 20mM (p ≤ 0.05) (Fig 2B). In parallel, metformin increased the swimming orientation persistence time (τ_p_), indicating that bacteria maintained their swimming direction for longer periods before reorientation (Fig 2C). These results suggest an enhancement of persistence runs, allowing bacteria to move more efficiently along gradients, a result also consistent with the higher abundance of *cheZ* and lower abundance of *cheA* found in the metagenomic analysis.

To determine further whether metformin influences directional migration in presence of chemical gradients, we exposed WT *E. coli* bacteria to a metformin gradient using a microfluidic system and tracked individual bacteria for long tracking times (*N* = 51 tracks for *T* = 120 s). The tracks show a manifest asymmetric spatial distribution, with cells preferentially moving to regions containing lower metformin concentrations (Fig 2D). Quantitative analysis of the chemotactic velocity drift in the bulk yields U_chembulk_ = -2.32 ± 0.70 μm/s, with a mean mode speed of 29.06 ± 0.94 μm/s. Most interestingly, this directional bias was not observed in the Δ*cheY* mutant which lacks a central component of the chemotaxis signaling pathway (Fig 2E, Fig S4). Bacteria would remain essentially on the upper or the lower surfaces performing circles ^52^, exhibiting no detectable directional bias along the metformin gradient (AD63 strain, U_chem_ = 0.24 μm/s ± 0.49, mean mode speed equal to 34.1 ± 1.2 μm/s) (Fig 2E, Fig S4).

Furthermore, compared to the WT AD62 strain, the Δ*cheY* mutant AD63 displayed greater sensitivity to metformin, with significant growth inhibition observed at concentrations as low as 1 mM, compared to 10 mM in AD62 (Fig S3). These findings indicate that an intact chemotaxis pathway (e.., CheY-dependent signaling) is required for the directional response to metformin and influences bacterial sensitivity to metformin exposure ^53,54^.

The directional bacterial response to metformin was further confirmed using soft-agar assays (Fig 2F-G). Similar directional responses were observed in other motile bacterial species, including the Gram-negative Proteobacteria *Pseudomonas aeruginosa* and *Salmonella Typhimurium*. These bacteria are frequently involved in gastrointestinal infections, particularly in cases of dysbiosis ^55,56^. We made the same observation with the Gram-positive species *Bacillus subtilis* and *Listeria monocytogenes*, although the magnitude of the response varied across species (Fig 2F–G, Fig S4). Notably, the concentrations associated with directional redistribution broadly overlapped with those affecting bacterial growth in each species (Fig 2F–G, Fig S3, Fig S4), indicating that sensitivity to metformin differs among bacterial taxa.

### The chemorepellent response to metformin required the chemotaxis pathway and tsr

To further investigate the molecular basis of metformin gradient detection, we used the Keio collection of single-gene deletion mutants to explore the response to metformin in individually deleted chemosensor genes ^57,58^. We focused on genes found enriched in our metagenomic analyses (Fig. 1C). Deletion of *trg* or *tar* did not abolish the chemorepellent response to metformin, indicating that these receptors are individually dispensable for metformin sensing under the condition tested. In contrast, deletion of *tsr* completely inhibited the directional migration away from the metformin gradient while basal motility was preserved on Brain Heart Infusion medium (BHI) alone. (Fig. 2H-I, Fig. S5). These results identify Tsr as the primary chemoreceptor required for the chemorepellent response to metformin ^59,60^. Deletion of *aer* partially attenuated the chemorepellent response in approximately half of the tested conditions, suggesting functional contribution of other chemosensors in the response (Fig. S5). Finally, deletion of *tap* abolished motility under both conditions, precluding any conclusion regarding its specific role in metformin sensing (Fig. S5). Interestingly, deletion of *tar* also resulted in the absence of run behavior on BHI alone, suggesting that Tar may be required for chemotaxis toward attractants present in this nutrient-rich medium — consistent with its well-established role as the primary receptor for aspartate and maltose ^61^. As a consequence, whether Tar plays an additional role in the response to metformin cannot be determined from the present experimental design, since the absence of directed motility in BHI alone precludes interpretation of the BHI + metformin condition.

Together, these results show that metformin acts as a bacterial chemorepellent through an intact chemotaxis pathway and provide experimental support for the altered chemotaxis and mobile gene signature observed in the human metagenomic analysis. This prompted us to investigate whether these behavioral changes potentially influence bacterial interactions with the mucus barrier ^55,56^.

### Metformin limits bacterial penetration into intestinal mucus

We next tested whether metformin-induced changes in bacterial behavior may affect interactions with the mucus layer using a previously described microfluidic platform using either mucin or mucus (Fig 3A) ^62^.

**Fig 3:**
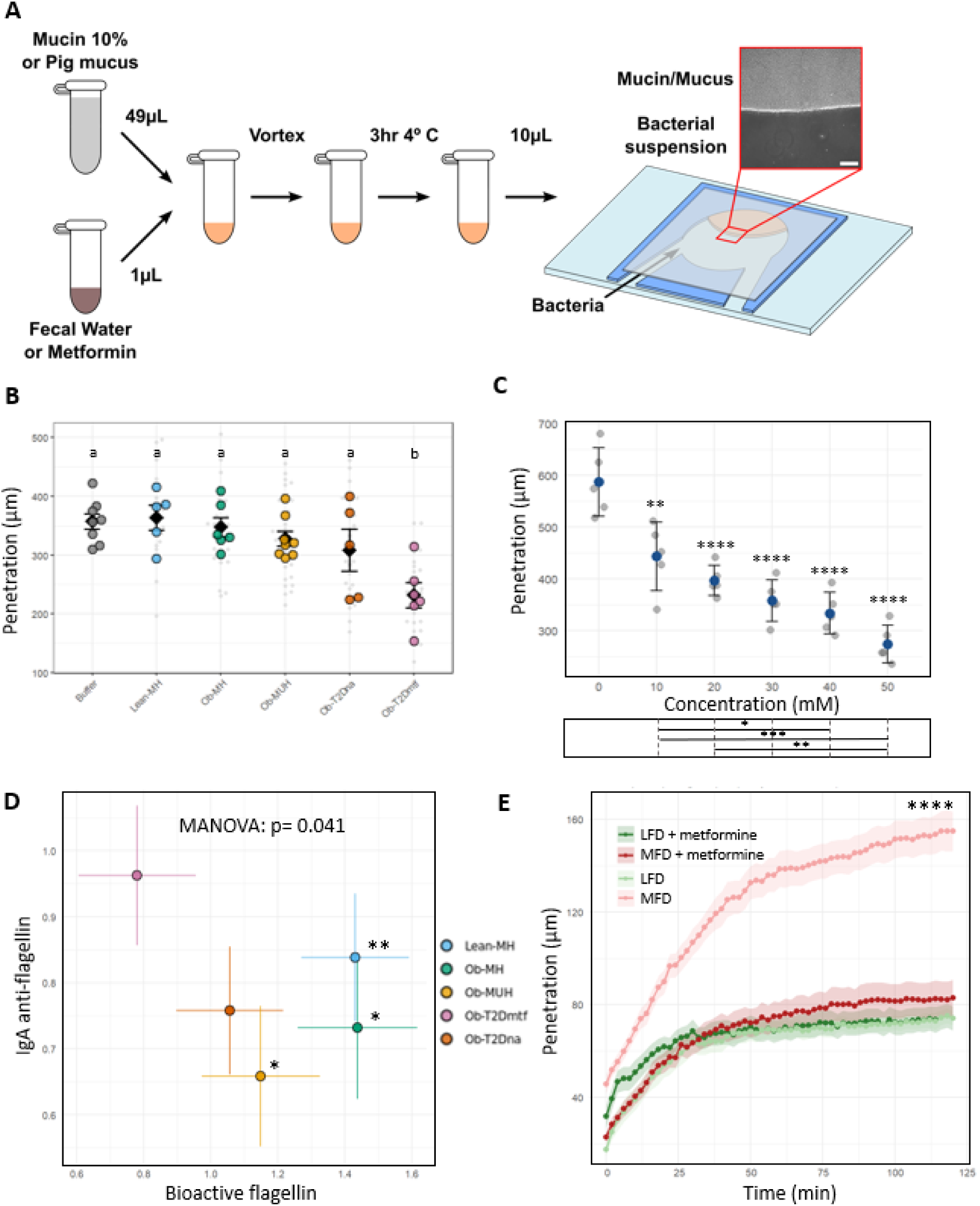
Metformin limits bacterial penetration into host mucus. **A:** Schematic diagram of the protocol and the microfluidic system used to monitor bacterial penetration into mucus or mucin in the presence of fecal water or metformin**. B.** Penetration distance of *E. coli* AD62 into mucin in the presence of fecal water from patients in the different study groups. Grey dots represent individual experimental replicates and colored dots represent biological coefficient of variation (CV)-based outlier filtering. A CV threshold of 20% (or 30% when necessary to limit data loss) was applied within each biological replicate; corresponding CV distributions and retained replicate numbers are provided in Fig S6 and Table S6. Statistical analyses were performed using a linear mixed-effects model (lmer) following assessment of normality by the Shapiro–Wilk test and BH correction. Group*s* sharing a letter are not significantly different; different letters indicate FDR ≤ 0.05. **C:** Penetration distance of *E. coli* through a mucin matrix was quantified after 120 min in the presence of the indicated metformin concentrations. Each point represents an independent experiment and bars indicate the mean ± SD. After assessment of normality by the Shapiro–Wilk test, statistical significance was assessed by one-way ANOVA followed by Tukey’s post-hoc test. Tukey-adjusted p-values for comparisons against the 0 mM condition are shown above the corresponding data points; Tukey-adjusted p-values for all pairwise comparisons between concentrations are reported in the table below. **D:** Estimated marginal means (EMM) of fecal anti-flagellin IgA levels and flagellin bioactivity, measured using a TLR5 reporter assay, adjusted for age and sex, across the different groups. Differences between groups were first assessed using a multivariate analysis of variance (MANOVA) based on Pillai’s trace statistic. Pairwise group comparisons were subsequently performed using Hotelling’s T² tests, and BH correction. Asterisks indicate groups that differ significantly from the Ob-T2Dmtf group. **E:** Temporal evolution of the penetration length of *E. coli* into mucus from the small intestine of pigs fed with low (n = 5, LFD) or medium fat diet (n = 5, MFD), in the presence or absence of metformin 50 mM in the mucus. Graphs represent the mean of all biological replicates per condition ± standard error of the mean (SEM). Each biological replicate was measured in technical triplicate. Statistical significance was assessed using a linear mixed-effects model (Time × Diet × Metformin, with biological replicate as a random effect), followed by pairwise comparisons between groups, with adjustment by Tukey method. Asterisks indicate a statistically significant difference between the MFD group and all other groups. No significant differences were observed among the remaining three groups (LFD, LFD+metformin, MFD+metformin).*: FDR ≤ 0.05, **: FDR ≤ 0.01, ****: FDR / Tukey-adjusted p-value≤ 0.000 ≤ 0.0001.

We first assessed bacterial penetration in mucin in the presence of fecal waters obtained from individuals belonging to the different study groups (N = 30) (Fig 3B). Among all groups, only fecal water from metformin-treated individuals (Ob-T2Dmtf) significantly reduced bacterial penetration into the mucin matrix compared with the other conditions (FDR ≤ 0.05) (Fig 3B, Fig S6, Table S6). No significant differences in mucin penetration were detected between Lean-MH individuals and the other metabolic groups in the absence of metformin treatment (Fig 3B). Importantly, direct exposure of mucin to metformin similarly induced a concentration-dependent reduction in *E. coli* penetration through the mucin matrix (Fig 3C, Table S7; and Supplementary Videos).

While the reduction of bacterial penetration in mucin, observed with fecal waters from metformin-treated individuals, suggests that luminal factors contribute to this phenotype, the intestinal environment is considerably more complex than the simplified *in vitro* systems. We therefore investigated whether metformin treatment could also be associated with changes in some host- and/or microbe-derived factors, previously associated with bacterial access to mucus. Based on a previously reported observation that IgA restricts bacterial penetration through intestinal mucus, we quantified anti-flagellin IgA in fecal waters from the patient groups, as well as bioactive flagellin, a major determinant of bacterial motility and mucus penetration ^62,63^. This was performed using the TLR5 reporter assay described above (Fig 2A). Fecal waters from metformin-treated individuals displayed significantly lower flagellin activity, together with higher levels of anti-flagellin IgA, as compared to those of the other groups (Fig 3D). This increase in anti-flagellin IgA is particularly relevant given that, in our *in vitro* mucin penetration assay, IgA reduced *E. coli* penetration through mucin more strongly than metformin alone, with the combination of both showing a further, albeit non-significant, decrease (Fig S7, Table S8). This suggests that the elevated IgA levels observed in metformin-treated individuals could act in concert with metformin’s direct effect on bacteria, potentially reinforcing bacterial exclusion from the mucus layer in vivo beyond what either factor achieves alone. Noteworthy, metagenomic analyses revealed a lower abundance of *fliC*, the gene encoding flagellin, in metformin-treated individuals (Fig 1C). These observations made on human fecal water is then in contrast with the *in vitro* finding that metformin direct exposure increased bioactive flagellin production in *E. coli* and highlight differences between bacterial responses measured in isolated experimental systems and those observed within some elements of the intestinal ecosystem (Fig 1C, 2A).

Second, we extended these observations in a more physiologically relevant setting and assessed bacterial penetration using *ex vivo* small-intestinal mucus isolated from pigs. This model more closely recapitulates the structural and biochemical properties of native intestinal mucus and overcomes the limited accessibility of fresh human intestinal mucus for functional studies. As previously described by Ranson et al., donor pigs were fed with either a low-fat or a moderate-fat diet, conditions known to modify mucus properties ^7,64^. Mucus from pigs fed a moderate-fat diet permitted greater *E. coli* penetration than mucus from low-fat-fed animals (Fig 3E, Fig 4, Table S9). Addition of metformin significantly reduced bacterial penetration in the moderate-fat condition, restoring values toward those observed in the low-fat group. In contrast, metformin did not reduce bacterial penetration into mucus from low-fat-fed animals. Thus, the effect of metformin was more apparent under conditions associated with greater baseline penetration (e.g., mucus exposed to increased fat consumption of pigs) (Fig 3E, Table S9).

**Fig 4:**
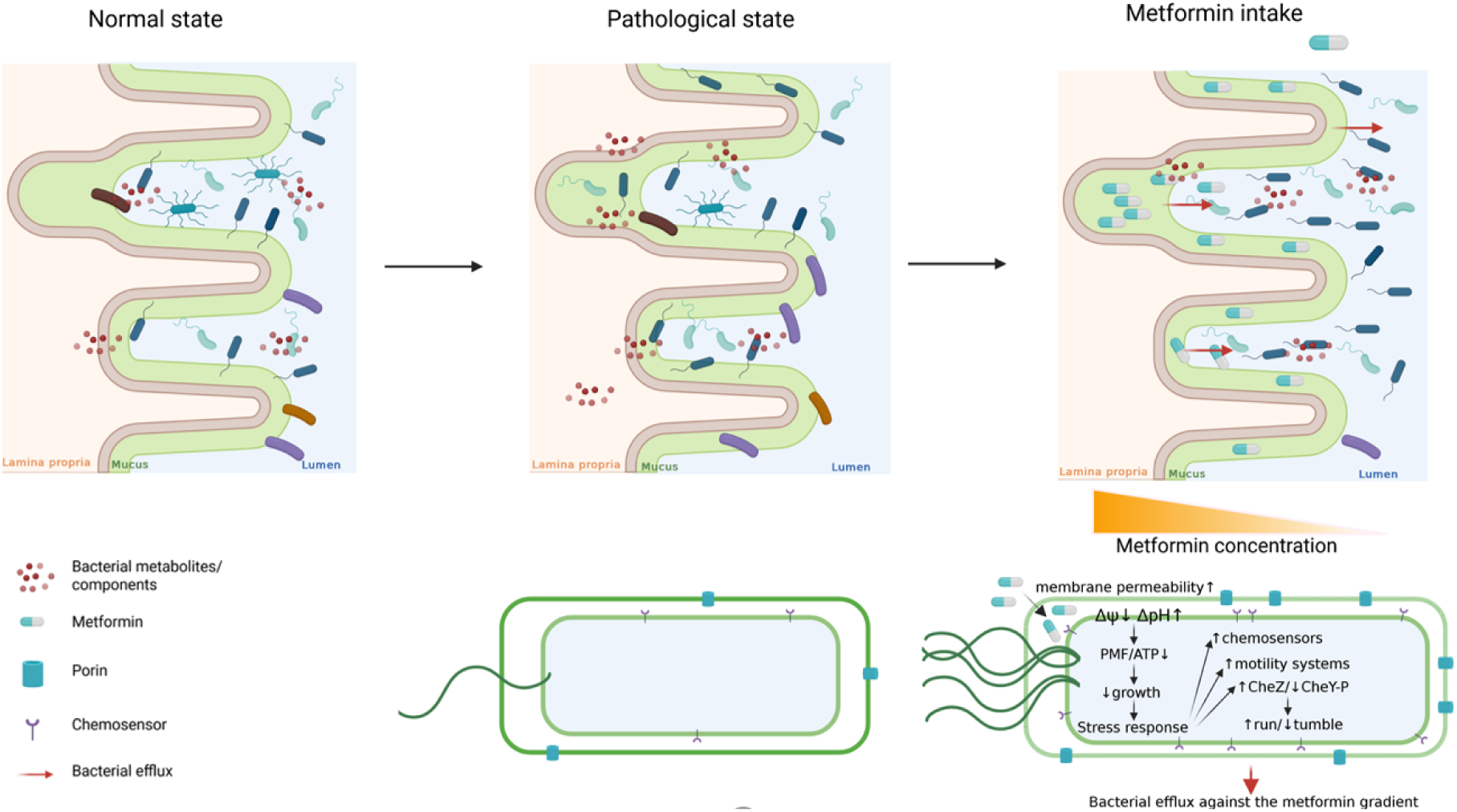
Proposed model by which metformin alters bacterial chemotaxis and limits penetration into intestinal mucus. Under homeostatic conditions, the intestinal mucus layer contribute to spatial segregation between the gut microbiota and the host epithelium. Conditions associated with altered mucus properties can increase bacterial penetration into the mucus layer and thereby enhance host exposure to microbial products. Following oral administration, metformin (a small, positively charged molecule) reaches millimolar concentration within the intestine compartment exposing resident bacteria to significant concentrations. Our findings support a model in which metformin influences both microbial community structure and bacterial behavior. Metformin treatment is associated with reduced microbial richness and increased abundance of some motile bacteria such as Proteobacteria, as well as with enrichment of genes involved in chemotaxis and flagellar mobility.

Together, these findings extend the observations obtained in the microfluidic mucin system and indicate that metformin limits bacterial penetration across distinct mucus environments, including conditions associated with increased bacterial access to the mucus layer.

At the bacterial level, metformin at millimolar concentration increases swimming speed and directional persistence and acts as a chemorepellent. This directional response requires an intact chemotaxis pathway and is abolished in Δ*cheY* bacteria and depends on the chemoreceptor Tsr, with a partial contribution from Aer.

Previous studies indicate that, at high concentrations, metformin can perturb bacterial membrane properties and bioenergetics, including transmembrane potential and proton motive force ^48,49^. Such perturbations may provide signals sensed through chemotaxis and energy-taxis pathways, although the precise upstream mechanism remains to be established.

Functionally, metformin reduces bacterial penetration into purified mucin and into native intestinal mucus under conditions associated with increased baseline penetration. Fecal water from metformin-treated individuals similarly restricts bacterial penetration and is characterized by reduced flagellin bioactivity and increased anti-flagellin IgA. Together, these bacterial and host-associated effects may contribute to maintaining microbiota–host spatial segregation at the intestinal mucus interface.

## Discussion

The main finding of this study is that metformin can act as a bacterial chemorepellent, reshaping bacterial behavior and modifying microbiota–host spatial segregation. By integrating human metagenomics with quantitative bacterial biophysics, we herein report for the first time that metformin-associated microbial signatures in humans translate into measurable changes in bacterial behavior and ultimately into altered interactions with the mucus barrier. We identify bacterial behavior as a functional dimension of the gut microbiome that cannot be inferred from the taxonomic composition alone.

Our findings help resolve an apparent paradox. Metformin treatment was associated with enrichment of chemotaxis- and motility-related functions in the gut microbiome, despite restricting bacterial penetration into mucin and intestinal mucus in our experimental systems. Metagenomic signatures were consistent with increased directional persistence, a prediction supported by single-cell tracking experiments showing that metformin altered bacterial navigation and induced a chemorepellent response in the tested concentration. These observations indicate that enrichment of microbial motility functions does not necessarily translate into increased bacterial access to the host.

At millimolar concentration, compatible with those potentially reached locally within the intestinal compartment ^24,25,65^, metformin acted as a chemorepellent, redirecting bacterial migration away from metformin gradients and thereby limiting penetration into both purified mucin and *ex vivo* intestinal mucus (Fig 4). Importantly, this effect was observed in porcine mucus from animals fed a moderate-fat diet, where bacterial penetration returned to levels comparable to those observed in low-fat fed animals. Although the underlying alterations in mucus composition remain to be defined and access to human mucus is difficult, these findings suggest that metformin may preferentially reinforce microbiota–host spatial segregation in altered mucus environments. Our observation is particularly relevant in light of accumulating evidence linking defective microbiota–host spatial segregation with obesity, insulin resistance and inflammatory diseases, where bacterial encroachment into the mucus layer may increase host exposure to microbial-associated molecular patterns ^11,63,66^. Thus, we propose a model in which metformin modifies microbiota–host spatial segregation by establishing concentration gradients within and outside the mucus layer that may limit effective bacterial access to the epithelial surface.

The mechanisms by which metformin produces this phenotype remain incompletely understood; nevertheless, our findings raise the possibility that metformin-induced perturbations of bacterial energetics are sensed through conserved chemotaxis and energy-sensing pathways, thereby translating drug exposure into directed bacterial escape from metformin-rich environments in our tested conditions ^48,49^. Behaviorally, this response was characterized by increased directional persistence, with extended run duration and reduced tumbling frequency. Mechanistically, this response requires an intact chemotaxis signaling cascade. Tsr, a methyl-accepting chemotaxis protein known to participate in sensing changes in bacterial energetic state, emerged as a major determinant of the response. Downstream, modulation of CheA activity and cheY phosphorylation controls flagellar motor switching, with reduced CheY-P favoring extended runs and directional persistence ^59,60^. Consistently, the response was abolished in the Δ*cheY* mutant, confirming that CheY-dependent motor switching is required for the metformin-induced behavioral response. Among the chemoreceptors tested, deletion of Tsr completely abolished directed motility away from metformin gradients, whereas deletion of Aer partially attenuated the response under a subset of conditions. Given the respective roles of Tsr and Aer in sensing interconnected aspects of bacterial energetic state ^59,60^, these findings raise the possibility that metformin-induced perturbations of bacterial bioenergetics contribute to the observed behavioral response. The underlying signal-including potential contributions of proton motive force and respiratory redox state-remains to be established. In contrast, deletion of Tar or Trg did not abrogate the response, indicating that these receptors are individually dispensable under the conditions tested. Moreover, similar chemorepellent responses were observed across phylogenetically distant Gram-positive and Gram-negative bacteria, suggesting that the capacity to respond to metformin may extend beyond a single bacterial group, including enteric bacteria with pathogenic potential.

Because metformin accumulates within the intestinal lumen and mucosa at concentrations far exceeding those detected in plasma, resident microorganisms are exposed to millimolar drug levels. ^24,25,65^. Studies have shown that metformin can perturb bacterial membrane function and cellular energetics, including processes linked to proton motive force and transmembrane electrochemical gradients (Δψ), and that these perturbations can be detected through conserved energy taxis systems involving Aer and MCPs, which couple intracellular energetic status to directional movement ^46–49,67–69^. Noteworthy, in our metagenomics observations, enrichment of MCPs and *cheZ,* together with depletion of *cheA,* is compatible with reduced CheY phosphorylation, suggesting fewer tumbling events and increased directional persistence, that matches with the behavioral phenotype we observed. Moreover, depletion of type IV pilus genes with enrichment of flagellar-associated functions may reflect altered motility in response to metformin exposure ^70,71^. These observations cannot establish causality or distinguish changes in bacterial behavior from shifts in community composition but they appear consistent with the phenotype observed across the different experimental systems. Metformin concentrations eliciting chemorepellent responses overlapped with those affecting bacterial growth, raising the possibility that the behavioral response is linked to metformin-induced changes in bacterial physiology and energetic state. Consistent with this interpretation, the Δ*fliC* mutant was less susceptible to metformin-induced growth inhibition than the WT, raising the possibility that activation of motility programs represents an energetically costly adaptive response to metformin exposure ^72,73^.

Because bacterial motility and spatial organization determine exposure of the host to immunostimulatory microbial ligands, altered bacterial navigation would be expected to influence mucosal immunity. Although metformin increased bioactive flagellin production in *E. coli* under *in vitro* conditions, fecal waters from metformin-treated individuals displayed lower flagellin bioactivity, together with increased anti-flagellin IgA levels. These observations indicate that direct bacterial responses to metformin occur within a host-regulated ecosystem in which immune mechanisms contribute to microbial containment. Previous studies have shown that metformin enhances intestinal IgA responses, while anti-flagellin IgA limits bacterial encroachment into mucus, consistent with the mechanism recently demonstrated by Tran et al. in mice ^63,74,75^. These findings suggest that metformin promotes microbiota–host spatial segregation through complementary bacterial and host mechanisms. By limiting bacterial access to the mucus layer and reducing exposure to pro-inflammatory microbial ligands such as flagellin, lipopolysaccharide or peptidoglycan, metformin may attenuate mucosal immune activation ^21,34,35,76–78^. Although this model requires direct experimental validation, it provides a physiological framework linking bacterial behavior, mucosal barrier function and improved metabolic homeostasis.

Our study has some limitations. The metagenomic analyses infer functional potential rather than pathway activity and cannot distinguish adaptive bacterial responses from shifts in community composition. Although the principal mechanistic studies were performed in *E. coli*, similar chemorepellent responses across phylogenetically distinct bacteria suggest that the phenomenon extends beyond a single species. The chemorepellent phenotype was characterized at relatively high metformin concentrations; while such concentrations may be reached locally within the intestinal compartment following oral administration, the spatial and temporal exposure of bacteria within the human gut and mucus remains insufficiently characterized. Future *in vivo* studies will be required to establish the physiological contribution of metformin-induced changes in bacterial behavior to microbiota–host spatial organization and metabolic outcomes.

Collectively, our findings support a model in which metformin acts not only as a metabolic drug but also as a microbiota-directed ecological signal that modulates bacterial behavior. Our work identifies bacterial behavior as an additional functional layer of the gut microbiome that cannot be inferred from taxonomic composition or metagenomic functional potential alone. Notably, the chemorepellent response observed under our experimental conditions extended beyond commensal bacteria to several enteric pathogens, suggesting that metformin can engage conserved bacterial sensing and motility pathways across diverse bacterial species. By restricting bacterial access to the mucosal compartment, such behavioral responses may contribute to microbiota–host spatial segregation and potentially to host protection. More broadly, incorporating bacterial behavior and the local microenvironment into microbiome research may provide a more integrated understanding of host–microbiota interactions and open new perspectives for microbiota-directed therapeutic strategies.

## Material & methods

### Ethics and Human participants

Human samples and clinical data were obtained from the French subset of the MetaCardis cohort. The promoter is Assistance publique-hôpitaux de Paris and the study protocol were approved by the Comité de Protection des Personnes (CPP) Île-de-France III [number P121102]. All participants provided written informed consent, and the study was conducted in accordance with the Declaration of Helsinki. MetaCardis is registered at ClinicalTrials.gov (NCT02059538).

### Human groups

The global MetaCardis cohort includes extensive clinical, lifestyle, and nutritional data, as well as fecal metagenomic sequencing, from 2,214 participants recruited in clinical centers in France, Germany, and Denmark between 2013 and 2015. For the present study, only participants recruited in France were included ^39,79–81^, because extracted fecal waters were available. In this French arm, there were 684 participants. Subjects were classified into their metabolic condition.

The selected subjects were classified according to their body mass index (BMI) into two groups: lean participants (Lean-MH; BMI<25 kg/m², n=137) and participant with obesity (Ob; BMI>=30 kg/m²). Participants with obesity were further stratified according to metabolic status into three categories reflecting increasing metabolic impairment. The first category comprised participants with obesity but metabolically healthy (Ob-MH, n=112), defined by the absence of metabolic syndrome and T2D. The second category included subjects with obesity but metabolically unhealthy (Ob-MUH, n=167), defined by the presence of metabolic syndrome according to the International Diabetes Federation criteria, and without T2D ^82^. The third category comprised subjects with obesity and T2D (Ob-T2D), defined according to the American Diabetes Association criteria, and was further subdivided into participants treated with metformin (Ob-T2Dmtf, n=184) and those not treated with metformin (Ob-T2Dna, n=84) ^83^. Briefly, exclusion criteria for the MetaCardis cohort included age >75 years, antibiotic use within the three months preceding inclusion, acute or chronic inflammatory or infectious diseases (including hepatitis C virus, hepatitis B virus, and HIV infection), autoimmune disease, severe renal impairment (estimated glomerular filtration rate <50 mL·min⁻¹·1.73 m⁻²), drug or alcohol addiction, a history of abdominal cancer, abdominal radiation therapy, intestinal resection (except appendectomy), or organ transplantation with immunosuppressive therapy.

### Metagenomic analysis

To account for differences in sequencing depth across samples, reads were randomly subsampled to a uniform depth within each sequencing batch using seqtk. The subsampling depth was defined by the sample with the lowest number of read pairs after exclusion of samples with insufficient sequencing depth, as assessed by inspection of rarefaction curves. Metagenomic analyses were performed using the bioBakery workflow. Read-level quality control was carried out using KneadData with default parameters, including quality filtering with Trimmomatic (SLIDINGWINDOW:4:20, MINLEN:75) and removal of human contaminant sequences by alignment to the human reference genome using Bowtie2 in very-sensitive mode. Taxonomic profiling was performed using MetaPhlAn4 with the mpa_vJan21_CHOCOPhlAnSGB_202103 reference database, which comprises 21,978 species-level genome bins (SGBs) derived from a reference gene catalog of 5.1 million taxonomic markers. Functional profiling was performed using HUMAnN3, which quantifies microbial gene family and metabolic pathway abundances from metagenomic data. Gene family abundances were normalized to copies per million reads (CPM) ^84,85^.

### Fecal bioactive flagellin

Bioactive flagellin was quantified as previously described using embryonic kidney (HEK)-Blue-mTLR5 cells (Invivogen, hkb-mtlr5). Fecal material was resuspended in PBS to a final 100’mg/mL concentration and homogenized without beads to avoid bacteria disruption. Samples were centrifuged twice at 8,000’×’g for 10’min at 4’°C, serially diluted the supernatant (1:25-1:3,125), and applied to mammalian cells. Purified *S. typhimurium*-derived flagellin (Invivogen) was used for standard curve determination using HEK-Blue-mTLR5. After 24’h of stimulation, cell culture supernatants were applied to the QUANTI-Blue medium (Invivogen, rep-qb1) to measure alkaline phosphatase activity at 620’nm after 30’min. Optical densities were transformed with the flagellin standard curve data and the dilution factor.

### Fecal anti-flagellin IgA

Sample preparation for immunoglobulin quantification has been previously described ^34,53^. Fecal material was resuspended to a final concentration of 33’mg/mL in a collection media consisting of 0.05’mg soybean trypsin inhibitor per ml of a 3:1 mixture of 1× PBS and 0.1’M EDTA. Following centrifugation at 1,800 rpm for 10’min, the supernatant was centrifuged again at 10,000’rpm for 10’min at 4’°C. Final supernatant was collected and stored with glycerol and 2’mM PMSF (Sigma, P-7626) at −’20’°C until analysis. The quantification of anti-flagellin-IgA has been previously described ^34^. In brief, 96-well half-area microtiter plates (Costar, Corning, New York) were coated with 50’ng/well of either *S. Typhimurium* flagellin in 9.6’pH bicarbonate buffer overnight at 4’°C. Supernatants were then applied at a 1:2 dilution for 1’h at 37’°C. After incubation and washing, the wells were incubated with horseradish peroxidase-linked anti-human IgA (SeraCare, 5220-0360). Quantification was then developed by adding 3,3’,5,5’-TMB, and the optical density was the difference between readings at 450’nm and 540’nm. The Ratio Bioactive Flagellin/Anti-Flagellin IgA parameter was calculated using the final data from each analysis.

### Mucin purification

The protocol for mucin purification was adapted from Miller and Hoskins (1981) ^86^. Briefly, 10 g of mucin from porcine stomach (Sigma-Aldrich), 2.92 g of NaCl and 1.58 g of Na_2_HPO_4_ were dissolved in 500 ml of cold Milli-Q water. The suspension was stirred for 15 min using a magnetic stirrer, and the pH was adjusted to 7.2 with NaOH. The mucin solution was then stirred for approximately 20 h at 4 °C. Next, the solution was transferred to 50 mL tubes and centrifuged at 3,200 g for 40 min at 4 °C. Following centrifugation, 20 mL of the supernatant was transferred to new 50 mL tubes containing 30 mL of ethanol pre-cooled to -20 °C. The solution was mixed by gently inverting the tubes five times and centrifuged at 3,200 g for 10 min at 4 C. The supernatant was discarded, and each pellet was resuspended in 6.7 mL of cold 0.1 M NaCl. Every three suspensions were combined into a single tube and 30 mL of ethanol pre-cooled at -20 °C was added to each tube. The solution was mixed by gently inverting the tubes 3 times, and then centrifuged at 3,200 g for 20 min at 4 °C. The supernatant was discarded and each pellet was resuspended in 20 mL of cold 0.1 M NaCl. 30 mL of ethanol pre-cooled to -20 °C was added to each tube, the solution was mixed by gently inverting the tubes 3 times, and then centrifuged at 3200 g for 20 min at 4 °C. The supernatants were discarded and each pellet was resuspended in 10 mL of cold Milli-Q water. The content of the tubes was transferred to a dialysis tubing (12–14 kDa molecular weight cut-off) and the mucin solution was dialyzed against Milli-Q water overnight at 4 °C with gentle stirring. The following day, the dialysis water was replaced every 4 h, and the final purified mucin solution was then collected and stored at - 20 °C until lyophilization for approximately 24 h. Lyophilization was performed in the laboratory of Nathalie Kapel (Medical School Department). The resulting mucin powder was stored at 4 °C until reconstitution in 1 mM Phosphate Buffer at a final concentration of 10% w/v.

### Pig mucus

Pig mucus is derived from the duodenal-jejunal region of the small intestine of pigs fed a diet with low or moderate fat content, described in Ranson et al. ^7^.

### Strains used

The tests were conducted using the following *E. coli* strains: AD62, AD63 (AD62 with the *cheY* gene deleted), MG1655, and MG1655 with the *fliC* gene deleted. The tests also included strains of *Bacillus subtilis* 168 trpC2, *Pseudomonas aeruginosa* AA2, *Salmonella typhimurium* SL1344 and *Listeria monocytogenes* 10403S. The strains deleted in chemosensors (*tap, tar, trg, tsr, aer*) come from the *E. coli* Keio Knockout Collection, containing a set of single-gene, knockout mutants for all nonessential genes in *E. coli* ^57,58^.

### Preparation of *E. coli* suspension

*E. coli* pre-cultures were started by inoculating an aliquot from a glycerol stock into in liquid Lysogeny broth (LB) (VWR Life Science) supplemented with 100 μg/mL ampicillin and cultured overnight at 30 ‘C in a shaking incubator at 200 rpm. Next, 50 µL of the overnight culture was transferred into 5 mL of Tryptone Broth (TB, 10 g/L Tryptone, 5 g/L NaCl) supplemented with 100 μg/mL ampicillin and grown at 30 ‘C until an OD_600_ of 0.3-0.5 was reached. Cells were then harvested by centrifugation at 3,290 g for 5 minutes at room temperature (25 °C), after which the pellet was resuspended in 1 mL of Berg’s Motility Buffer X2 (BMB X2, 20 mM Phosphate Buffer (PB 100 mM: 61.5 mM K_2_HPO_4_ + 38.5 mM KH_2_PO_4_), 7.8 g/L NaCl, and 0.2 mM EDTA) supplemented with 25 g/L L-Serine (Sigma). The bacterial suspension was adjusted to an OD_600_ of 0.2 and finally mixed with and equal volume of Percoll (MP Biomedicals) to obtain a non-buoyant bacterial suspension and minimize cell sedimentation.

### Interface formation and Time-Lapse imaging

The formation of an interface between a bacterial suspension and mucus or mucin samples was performed using a previously described microfluidic setup ^62^. The device consists of a circular chamber 14 mm in diameter 200 μm deep, with an inlet and an outlet that allow the filling of the chamber by capillarity. Prior to each experiment, purified mucus or mucin samples were thawed for 1 h at room temperature on a tube rotator and homogenized by vortexing. For experiments with a mucin or mucus barrier, 10 μL of sample was deposited in one edge of the chamber, opposite to the edge that contains the inlet and outlet. The sample was carefully spread in order to create a straight interface. A glass coverslip was immediately placed over the chamber, which was then filled with the bacterial suspension prepared in a 1:1 mixture of BMBx2 and Percoll, allowing the suspension to slowly come into contact the mucus or mucin interface. After filling, the inlet and outlet were sealed with nail polish (Minute Quick-Finish Dry Varnish, Mavala) to minimize evaporation. Immediately after preparation, the microfluidic devices were transferred to an inverted epifluorescence microscope (Zeiss Observer Z1) equipped with a motorized stage and a 5X objective for automated time-lapse image acquisition at room temperature. Images (2,048×2,048 pixels) were acquired every 2 min using a Hamamatsu ORCA Fusion camera with illumination provided by a Zeiss Colibri 7 blue LED.

### Mucin/Mucus supplementation

For experiments in which mucin was supplemented with fecal water, 1 µL of fecal water was added to 49 µL of purified mucin 10% w/v. For experiments in which mucin was supplemented with metformin, 75 µL of purified mucin 10% w/v was mixed with 5 µL of metformin solution to obtain the desired final concentration in mucin. When required, immunoglobulin A (IgA) purified from human breast milk was added to the mucin to reach a final concentration of 0.077 µg/mL. The purified IgA was provided by D. Sterlin and G. Gorochov laboratory. Fresh breast milk of healthy lactating women was obtained from the Lactarium regional d’Ile de France (Hopital Necker Enfants-Malades) and approved by the ethical committee (Aves CENEM 2020-V4 Biocol lactoth’eque V1.0 20190408). Secretory IgA was purified using peptide M (InvivoGen) as described in Sterlin, D. et al., 2020 ^87^. For experiments in which mucus was supplemented with metformin, 80 µL of mucus was centrifuged at 3,290 g for 10 minutes, and 5 µL of the supernatant was replaced with 5 µL of metformin solution 0.8 M to obtain a final metformin concentration of 50 mM in the mucus. In all cases, samples were homogenized by vortexing and incubated for 1 h at room temperature prior to loading into the microfluidic device.

### Image analysis

Image processing and analysis were conducted using the Fiji distribution of ImageJ ^88^. Background was removed from each image prior to analysis using the “Subtract Background” function.

### Determination of the penetration length

For mucus samples, the penetration length was determined from the intensity profile of the images using the *getProfile* function in Fiji. The resulting profiles were fitted with an exponentially decaying function of the form y = Ae−λx + B, starting from the position of maximum intensity. The inverse of the decay constant, 1/λ, defines the penetration length λp, which represents the characteristic length scale over which the bacterial concentration decreases from its maximal value.

For mucin samples, bacterial cells were first detected using a Laplacian-of-Gaussian (LoG) spot detection approach implemented in custom Python code. Each image was filtered with a LoG operator and inverted to enhance bright, spot-like structures. Candidate cells were identified as local maxima using *peak_local_max* with a minimum separation constraint. Detections were then refined using subpixel localization and filtered based on the mean fluorescence intensity computed within a circular region corresponding to the expected size of a bacterium. The resulting cell positions were then projected onto the x-axis and binned to generate spatial histograms for each frame of the image stack. This spatial distribution was then fitted with the same exponentially decay function to compute the penetration length.

### *In vitro* bioactive flagellin

Bacteria in the exponential phase were diluted at OD 0.01 in media containing different concentrations of metformin (0, 10, 20, 50 mM) and incubated at 37°C for 5 hours. Bacterial cultures were then applied to mammalian cells. Purified *E. coli* flagellin (Invivogen) was used for standard curve determination using HEK-Blue-mTLR5. After 24’h of stimulation, cell culture supernatants were applied to the QUANTI-Blue medium (Invivogen, rep-qb1) to measure alkaline phosphatase activity at 620’nm after 30’min. Optical densities were transformed with the flagellin standard curve data and the dilution factor. The test was conducted in biological triplicate and in technical duplicate.

### Growth curve

The bacteria were diluted to an OD of 0.01 in LB from a culture in the exponential phase and then added to a 96-well flat-bottom plate (Sigma-Aldrich, Z707902-108EA) containing different concentrations of metformin (0 mM, 1 mM, 5 mM, 10 mM, 20 mM, 40 mM). The OD was monitored for 24 hours using a multimode microplate reader (TECAN, Spark) with readings taken every 15 minutes. The experiments were conducted using biological triplicates of technical duplicates.

### *E. coli* tracking

Single-cell tracking of *E. coli* was performed using a Lagrangian 3D tracking system as previously described ^50^. The approach enabled the extraction of individual cell motility parameters, including swimming velocity and orientation correlation. The orientation correlation time (τ), or persistence time, represents the characteristic timescale over which bacteria maintain their swimming direction; therefore, a higher τ indicates more persistent trajectories and less frequent reorientation events, consistent with an escape-like behavior. For the experiments conducted at different concentrations of metformin, bacterial suspensions were prepared by inoculating 50 µL of an overnight culture into 5 mL of TB medium supplemented with the appropriate concentration of metformin. Bacteria were grown for 3 h at 30 ‘C until reaching OD_600_ of 0.3-0.5. Cells were then harvested by centrifugation and resuspended in 1 mL of BMB X2 supplemented with 25 g/L L-Serine. The bacterial suspension was adjusted to an OD_600_ of 0.1, and 1 µL was diluted in 100 µL of a 1:1 mixture of BMB X2 (with L-Serine) and Percoll containing the appropriate concentration of metformin. The resulting suspension was introduced into a sealed chamber for single-cell tracking. The same preparation protocol was used for experiments performed in a metformin gradient, except that bacteria were exposed to metformin only after introduction into the microfluidic device. In both experimental configurations, the same Lagrangian 3D tracking system was used to quantify single-cell motility parameters.

For experiments performed at different metformin concentrations, bacterial trajectories were recorded at 80 fps and at room temperature (25°C) in three independent experiments performed on separate days. Tracks shorter than 45 s were excluded from the analysis. A total of 25, 39, and 38 bacterial tracks were analyzed for the 0, 10, and 20 mM metformin conditions, respectively. The mean tracking duration was 101.34 ± 42.62 s for 0 mM metformin, 99.81 ± 45.30 s for 10 mM metformin, and 140.33 ± 46.66 s for 20 mM metformin. Trajectories were smoothed using a Savitzky-Golay filter with a window of 0.2 s and analyzed using custom MATLAB code ^89^. Non-parametric, unpaired two-tailed Mann-Whitney tests for pairwise comparisons (α = 5%) were performed using GraphPad 6.01 Prism for Windows (GraphPad Software, http://www.graphpad.com).

For bacterial trajectories recorded in a metformin gradient, WT *E. coli* was exposed to a metformin gradient (50 mM/mm, spanning concentrations of 0–50 mM) generated using the commercial microfluidic device Ibidi® µ-Slide Chemotaxis cells (https://ibidi.com/channel-slides/9--slide-chemotaxis.html). Trajectories were also recorded at 80 fps at room temperature, and smoothed using a Savitzky-Golay filter with a window of 0.2 s. Three independent experiments were performed, yielding a total of 51 tracks of duration T = 120 s, ensuring that each bacterium carried the same statistical weight in the subsequent analysis. Gradient equilibration was estimated from the diffusion coefficients of metformin (D ≈ 1.23 × 10⁻⁵ cm²/s and fluorescein (D ≈ 0.43 × 10⁻⁵ cm²/s, used as a reference tracer ^90,91^. Since diffusive timescales scale inversely with D, and fluorescein diffuses roughly three times slower than metformin, a metformin gradient equilibrates faster and remains stable for about a third as long. As fluorescein gradients are stable from 30 min to at least 5 h post-injection (our observations, consistent with Grognot and Taute, 2021 ^92^), the metformin gradient is expected to remain stable for at least 100 min — the time window within which all experiments were performed. Bacteria were introduced simultaneously into both reservoirs to avoid diffusive drift. Each track was separated into bulk and surface events. Surface events were identified using a double-threshold criterion as defined in Junot et al., 2022 ^43^. The z-position distribution was fitted with a double Gaussian, whose maxima define the surface positions (z = 0 and z = H). Two thresholds, h1 = 8 μm and h2 = 3 μm, were set from each surface: a bacterium had to cross both consecutively to be assigned to a surface segment, and cross both again in reverse order to escape it; crossing h2 alone did not count as entry or exit. Surface residence time was defined as the interval between the first and last crossing of h2. Non-surface intervals were concatenated per bacterium into a bulk track, from which the chemotactic drift velocity was computed as Δx/Δt, where Δx is the total bulk displacement along x and Δt the total time spent in the bulk. All analyses were performed using home-made MATLAB code.

For each track, the director vectors pointing along the track are determined as 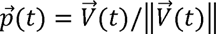. For each trajectory, we compute the autocorrelation function 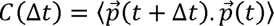. for a time lag Δt. The bracket denotes an average over a time window sliding along the track. To ensure good statistics, the maximum lag time Δt is chosen as one-tenth of the total track duration. The persistence time τ_p_ is obtained by fitting C(Δt) with an exponential decaying function 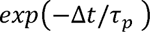.

### Metformin quantification by LC–MS/MS

Fecal material was resuspended in PBS to a final concentration of 100 mg mL⁻¹ and homogenized without beads to avoid bacterial disruption. Samples were centrifuged twice at 8,000 × g for 10 min at 4’°C. For metformin extraction, 150 μL of LC–MS grade acetonitrile was added to 50 μL of fecal water, vortexed for 10 s, incubated on ice for 15 min, and centrifuged at 15,600 × g for 10 min at 4’°C. Supernatants were evaporated to dryness using a vacuum concentrator (Concentrator Plus, Eppendorf) for 45 min at 30’°C (V-AL mode) and reconstituted in LC–MS grade water containing 10 mM ammonium formate.

Metformin was quantified using reversed-phase liquid chromatography coupled to high-resolution tandem mass spectrometry (LC–MS/MS, Thermo Scientific Q-Exactive Focus Hybrid Quadrupole-Orbitrap Mass Spectrometer). Chromatographic separation was performed on a Luna Omega PS C18 column (150 × 2.1 mm, 1.6 μm particle size; Phenomenex) maintained at 50’°C and operated at a flow rate of 0.4 mL min⁻¹. Mobile phase A consisted of 10 mM ammonium formate supplemented with 0.1% (v/v) formic acid in water, and mobile phase B consisted of acetonitrile containing 0.1% (v/v) formic acid. The gradient was as follows: 2% B from 0–1 min, increased to 20% B at 10 min, then to 95% B at 12 min, held at 95% B until 14 min, returned to 2% B at 14.1 min, and equilibrated until 16 min.

Mass spectrometric analyses were performed in positive electrospray ionization mode using full-scan acquisition (m/z 50–500, resolving power 70,000) combined with data-dependent MS/MS acquisition. Metformin was detected as the protonated molecular ion ([M+H]⁺, m/z 130.1087), and identity was confirmed by MS/MS fragmentation using an inclusion list targeting m/z 130.1087. MS/MS spectra were acquired at a resolving power of 17,500 with a 3.0 m/z isolation window and normalized collision energy ramped from 10 to 20. Characteristic fragment ions at m/z 71.0606 and 60.0556 were used to support metabolite identification.

External calibration standards of metformin (0.1 µM–20 mM) were prepared and analyzed under identical chromatographic and mass spectrometric conditions. Quantification was based on the extracted ion chromatogram of the protonated metformin ion.

### Metformin gradient agar

BHI agar plates were prepared with a metformin gradient ranging from 0 to 40 mM in a layer of 1 % agar, then covered with 0.3 % BHI agar without metformin. 10 µL of bacterial cultures in the exponential growth phase were spotted in the center of the plate. The experiments were performed in triplicate using a biological duplicate design. The same experiment was performed using 1 % BHI without metformin for the first layer.

### Microbiome analyses

To account for differences in sequencing depth, metagenomic reads were randomly subsampled to a defined number of read pairs per sample using seqtk. Alpha diversity was calculated from the subsampled data. Beta diversity was assessed using Bray–Curtis dissimilarities and visualized by principal coordinate analysis (PCoA). Differences in microbial community composition between groups were assessed by permutational multivariate analysis of variance (PERMANOVA) using the adonis2 function in the vegan package, with 99,999 permutations. Differential abundance of motility-related KEGG orthologues (KOs) was assessed using MaAsLin2 (version 1.26.0) with a linear model (analysis_method = “LM”). Clinical group, age and sex were included as fixed effects. No prevalence or abundance filtering was applied (min_prevalence = 0, min_abundance = 0). Abundance values were not normalized (normalization = “NONE”) and were log-transformed (transform = “LOG”) before modelling. P values were adjusted for multiple testing using the Benjamini–Hochberg (BH) procedure.

### Statistical analysis

Statistical analyses were performed using R (version 4.3.3). Where multiple hypotheses were tested, *P* values were adjusted using the BH procedure to control the false discovery rate (FDR). For post hoc pairwise comparisons following analysis of variance or mixed-effects models, Tukey adjustment was used to control the family-wise error rate. Exact statistical tests, sample sizes and definitions of replicates are provided in the corresponding figure legends.

A biological replicate was defined as an independent biological sample (an independent bacterial culture, animal or human participant, depending on the experiment), whereas technical replicates corresponded to repeated measurements of the same biological replicate. Technical replicates were either averaged before statistical analysis or explicitly accounted for as measurements nested within biological replicate using a random-effect term and were not treated as independent biological observations. Sample sizes for *in vitro* experiments were based on established experimental practice and biological reproducibility. Sample sizes for experiments using human or porcine material were constrained by sample availability. For analyses relying on parametric assumptions, normality was assessed using the Shapiro–Wilk test, applied to model residuals for ANOVA and mixed-effects models or to raw values where appropriate. When assumptions for parametric testing were not met, the corresponding non-parametric test was used. Detailed statistical analysis regarding each experiment are presented in supplemental information.

## Availability of data and materials

Raw sequencing human data used in this study are deposited in the ENA repository under accession codes PRJEB41311, PRJEB38742, and PRJEB37249 with public access. The phenotypical datasets of the current study are available from the MetaCardis cohort coordinator (KC) on reasonable request.

## Contributions

Conception & design: KS, AR, MVG, KC and EC. Performing experiments: KS, AR, MVG, HR, CD. Resources: MVG, KC, EC. Data analysis: KS, AR, ALG. Bioinformatic & statistical analyses: AR. Supervision: BC, RW, IGB, KC, EC. Writing-original draft: AR. Writing-review & editing: KS, AR, EC, KC. All authors have read and approved the current version of the manuscript.

## Acknowledgments

This work was supported by the Assistance Publique–Hôpitaux de Paris (AP-HP), sponsor of the MetaCardis clinical study, and by the European Union Seventh Framework Programme (FP7) through the MetaCardis consortium. The authors thanks French clinical investigators (nutrition, diabetology and cardiology department and the groups at Metagenopolis (INRAE, France), which performed metagenomic sequencing (Dr D. Ehrlich group). This work also received funding from the Marie Skłodowska-Curie Actions under Grant Agreement No. 101024357, the European Innovation Council (EIC) Pathfinder Challenge project NUTRIMMUNE (Grant Agreement No. 101162457) and well as the Food Systems, Microbiomes and Health Priority Research Programme and Equipment (PEPR SAMS) funded by the French Government through France 2030, managed by the Agence Nationale de la Recherche (ANR) (ANR-24-PESA-0010). Substantial support was provided by the ANR PushPull project (ANR-22-CE30-0038-04) and the ANR Bargain project focused on metformin effects on microbiome. Research in the BGPB team is supported by the Agence Nationale de la Recherche (ANR) through PEPR SAMS(ANR-24-PESA-0014), PPR AMR (ANR-20-PAMR-007) and ANR-24-CE17-1756. Research in the GG team is supported by the ANR through the BactRiA program (ANR-21-CE17-0037) and the PEPR TRANSCEND-ID program, by Inserm through the Impact Santé ANTIBAX Program and the BIOLIV MESSIDOR Grant 2024, the French Foundation for Medical Research (FRM EQU202203014622), and Région Ile de France through the DIM-ITAC program. Research in BC’s laboratory is supported by a Consolidator Grant (grant agreement InterBiome No. ERC-2024-CoG-101170920) from the European Research Council (ERC) under the European Union’s Horizon 2020 research and innovation program, ANR grants EMULBIONT (ANR-21-CE15-0042-01) and DREAM (ANR-20-PAMR-0002), région Ile-de-France DIM One Health-DOH 2.0, grant from the AFA Crohn RCH France and from the French government through the France 2030 investment plan managed by the National Research Agency (ANR), as part of the ANR-24-PESA-0008 and ANR 23 IAHU 0012. Axel Ranson is the recipient of a PhD fellowship from the École Normale Supérieure (ENS), Paris. The funders had no role in study design; data collection, analysis or interpretation; manuscript preparation; or the decision to submit the work for publication.

## Supplementary data

**Table S1:**
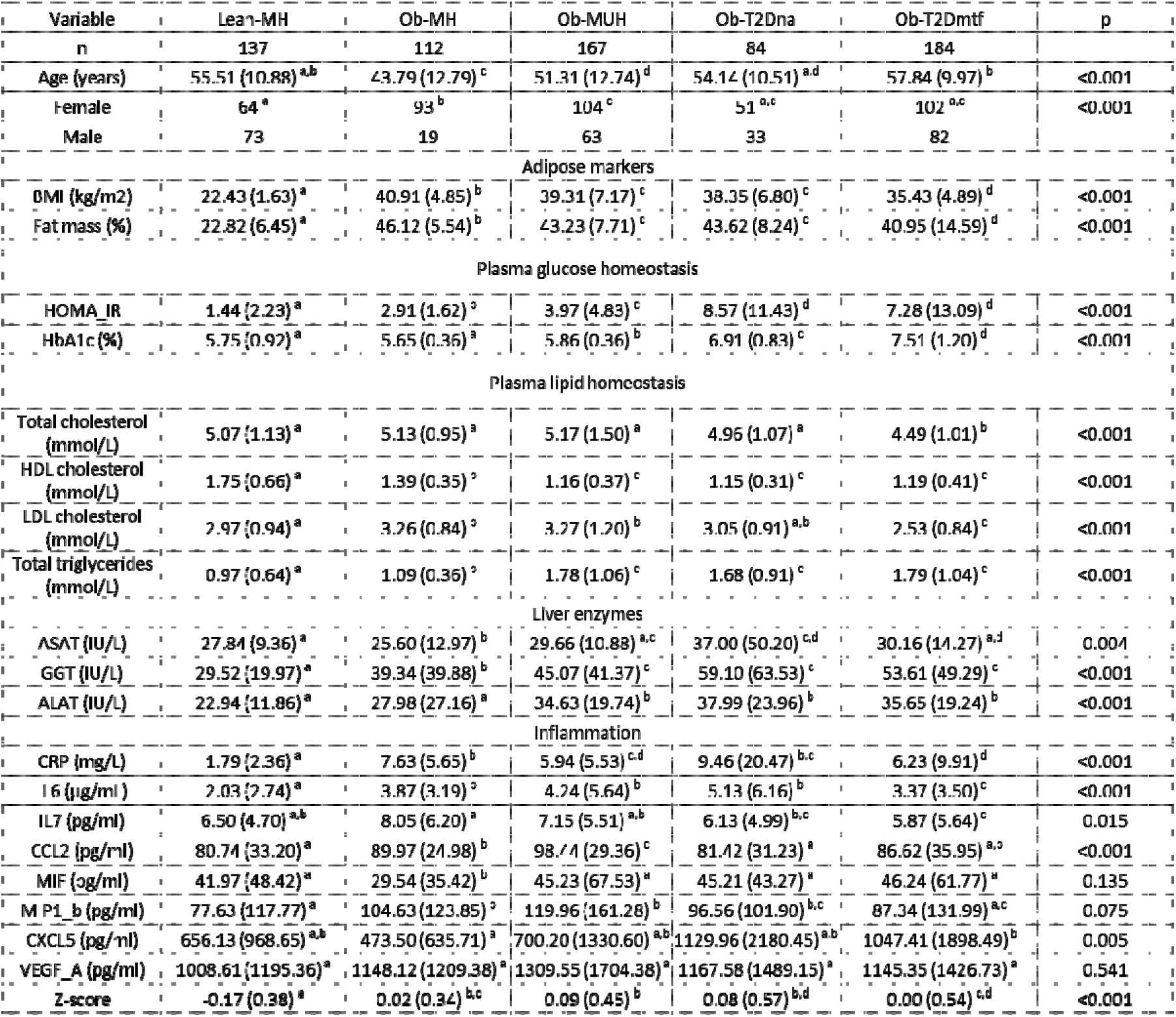
Clinical data on human French groups from MetaCardis. Data are shown as mean ± SD. P-values were calculated using the Wilcoxon rank-sum test for continuous variables and the χ² test for categorical variables (sex). Z-score (composite inflammation): a composite score calculated from all measured inflammatory markers. For each marker, individual values were standardized using the formula: Zi=(xi−μ)/ σ, where xi is the individual value, μ is the mean, and σ is the standard deviation of the entire cohort. The composite Z-score corresponds to the average of the individual Z-scores for each marker. A positive Z-score reflects an overall inflammatory profile above the cohort mean. Values are expressed as mean (standard deviation). For each variable, groups sharing the same superscript letter do not differ significantly, whereas groups with different letters are significantly different (p ≤ 0.05).

**Table S2:**
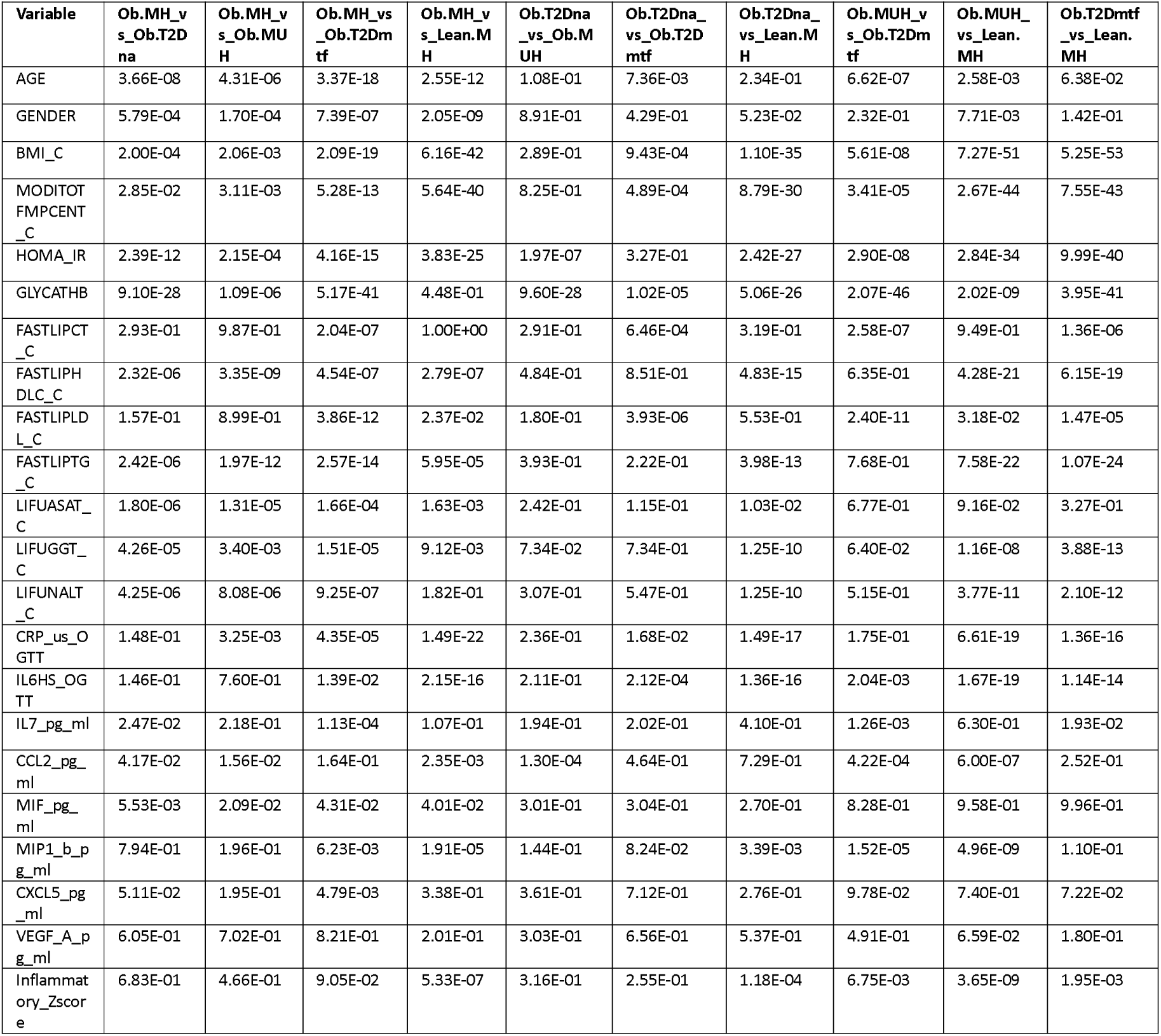
Statistical analyses of the clinical data on human cohort. Measured p-values between groups for the clinical variables presented in Table S1.

**Table S3: Statistical analyses of gut microbiota alpha and beta diversity across experimental groups, presented in Fig 1A and S1A-B**

*A:* Statistical comparisons of alpha diversity indices between experimental groups. The table reports the diversity index analyzed, the groups compared, pvalue, and BH-adjusted pvalue (FDR). *B:* Statistical analyses of beta diversity between experimental groups based on community composition. The table reports the distance metric used, test statistic, *R*², pvalue, and BH-adjusted pvalue (FDR).

**Table S4:** Keywords used for KO analyses.

|  |  |
| --- | --- |
| **Flagellar** | flagellar, flagellin |
| **Porin** | ompC, ompF |
| **Chemotaxis** | chemotaxis, aer |
| **Twitching** | twitching, type.IV.pilus, pilus.assembly,<br>prepilin.peptidase |
| **Gliding** | gliding |

**Table S5: Differential abundance analysis of motility-related genes across experimental groups using MaAsLin2 of Fig 1C**

Complete results of the multivariable association analysis performed with MaAsLin2 to identify differentially abundant motility-related genes between experimental groups. For each gene, the table reports the name of the gene and its category, the group studied, estimated coefficient, standard error, nominal Pvalue, BH-adjusted Pvalue (FDR) and the corresponding symbol and the number of samples in the group. Positive coefficients indicate higher relative abundance in the indicated group relative to the reference group, whereas negative coefficients indicate lower relative abundance.

**Fig S1:**
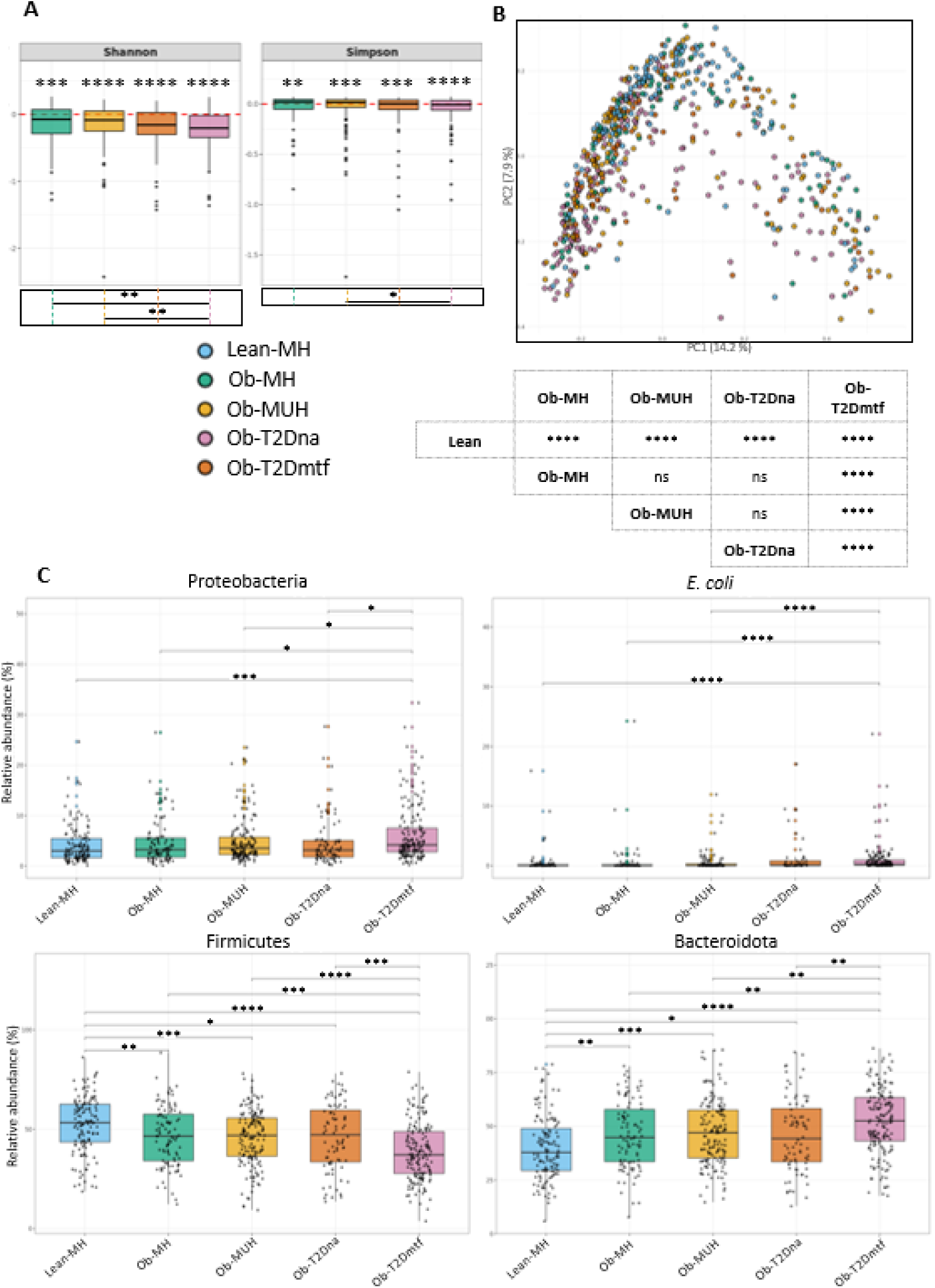
Metformin alters in gut microbiota structure and composition. **A**: Log-transformed fold change in alpha diversity (Shannon and Simpson) relative to the Lean-MH group. The dashed red line indicates the Lean-MH reference value (log fold change = 0). Pairwise comparisons were performed using Wilcoxon rank-sum tests, and p-values were adjusted for multiple testing using the BH procedure. Adjusted FDR values (***) above the graph correspond to comparisons of each clinical group to the Lean-MH group, whereas FDR values (***) in the box below correspond to comparisons between diseased groups. **B**: PCoA based on Bray-Curtis distance evaluating the bacterial composition of different groups with table with the statistic results. **C**: Analysis of the relative abundance of different taxonomic levels (Proteobacteria, *E. coli*, Firmicutes, Bacteroidota) across the different groups. Pairwise comparisons were performed using Wilcoxon rank-sum tests, and p-values were adjusted for multiple testing using the BH procedure. The absence of a statistical comparison between two groups indicates that the result is not significant. *: FDR ≤ 0.05, **: FDR ≤ 0.01, ***: FDR ≤ 0.001, ****: FDR ≤ 0.0001.

**Fig S2:**
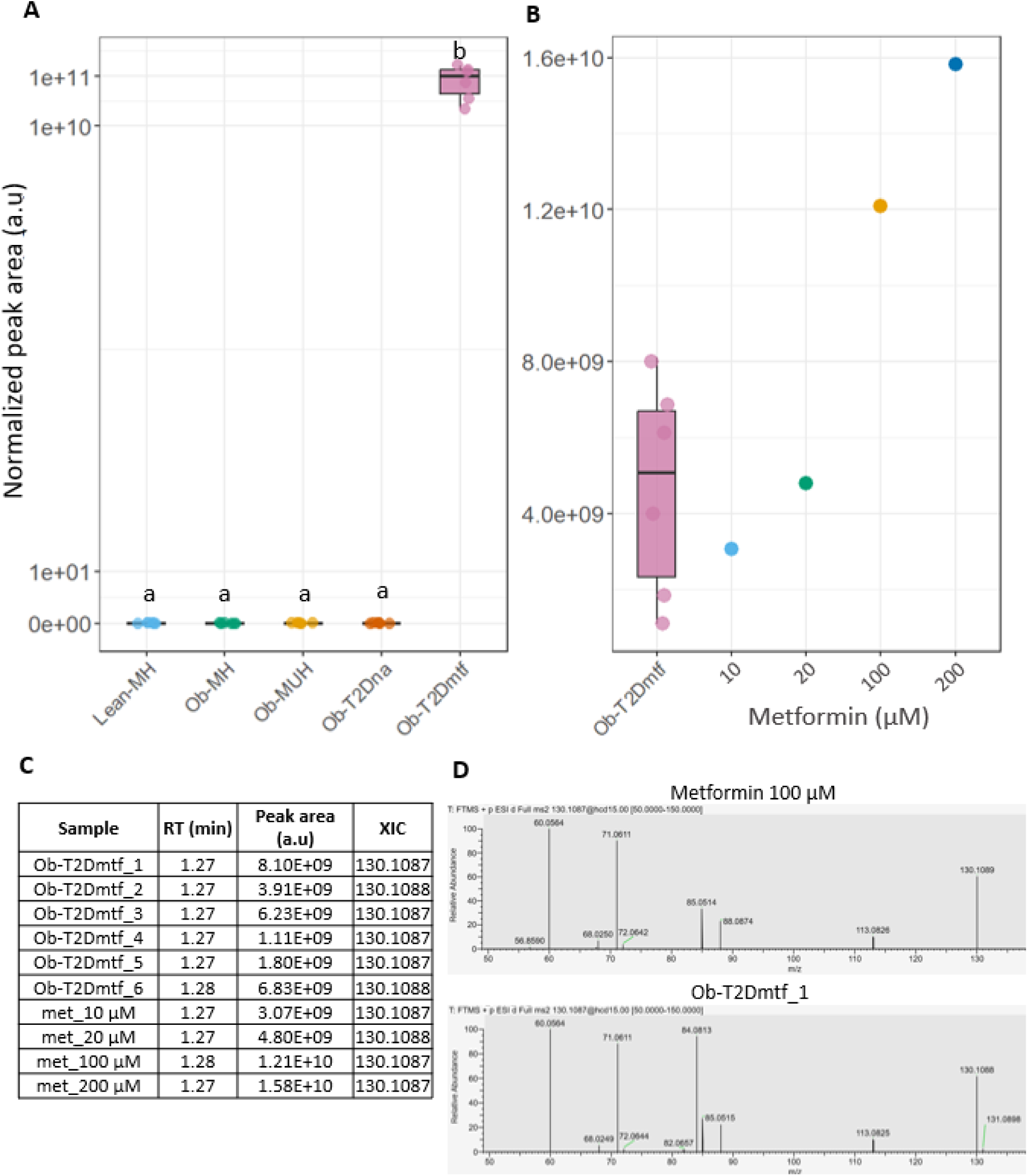
Fecal metformin levels across the different clinical groups measured by LC–MS/MS. **A:** Metformin was quantified in fecal water by LC–MS/MS across the different clinical groups. Metformin was detected in samples from metformin-treated individuals but not in the other groups. Fecal samples were diluted prior to LC-MS analysis (1:10 applied during fecal water preparation and 1:20 dilution prior to injection to prevent LC–MS saturation for Ob-T2Dmtf samples). Peak areas were corrected by subtracting the background signal measured in acetonitrile blank samples. Individual points represent biological replicates. Statistical differences between groups were assessed using pairwise Wilcoxon rank-sum tests followed by BH correction for multiple testing. Groups not sharing the same letter are significantly different (FDR ≤ 0.05). **B:** Quantification of the 200-fold diluted Ob-T2Dmtf samples, using a metformin standard curve, indicated metformin concentrations corresponding to approximately 1-10 mM in the stools, depending on the individual. **C:** Representative retention time, peak area and XIC of samples containing metformin. A signal at m/z 130.1088, corresponding to the theoretical m/z of protonated metformin ([M+H]⁺), was detected in Ob-T2Dmtf group with retention times similar to those of the control samples. *D*: Fragmentation profile of metformin ion of one standard (metformin 100 µM) and one Ob-T2Dmtf sample, used to support metabolite identification. The other standards and samples also exhibited the same characteristic fragment ions at m/z 71.0606 and 60.0556 were used. XIC: Extracted Ion Chromatogram; RT: Retention Time.

**Fig S3:**
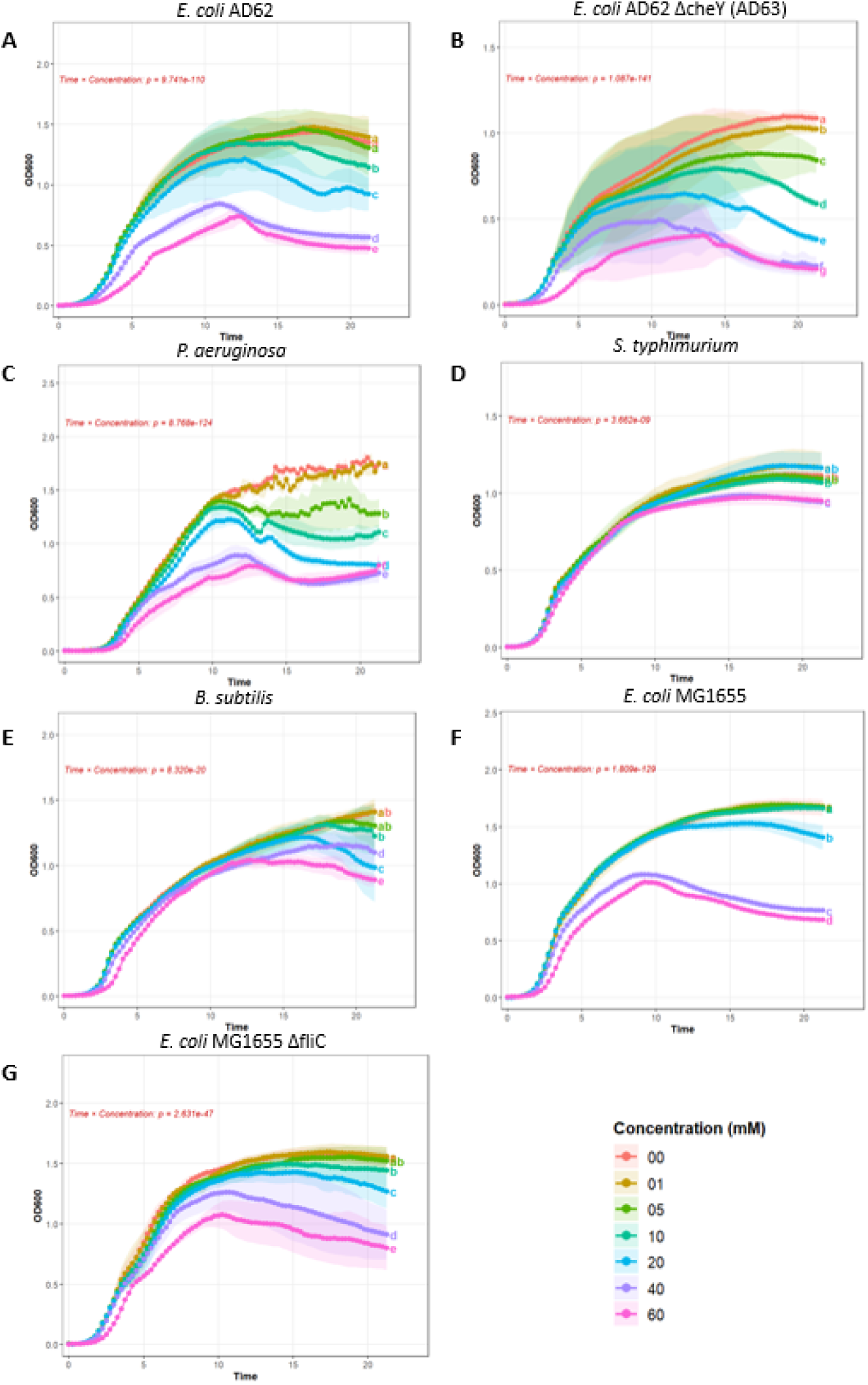
Growth kinetics of bacterial strains exposed to increasing metformin concentrations. Bacterial growth was monitored by measuring optical density (600nm) over time in the presence of the indicated metformin concentrations. Curves represent the mean ± SEM of 4 biological replicates in 3 technical replicates for E. coli AD62 (A), E. coli AD62 Δ*cheY* (AD63) (B), P. aeruginosa (C), *S. typhimurium* (D), *B. subtilis* (E), *E. coli* MG1655 (F) and *E. coli* MG1655 Δ*fliC* (G). Growth responses were analyzed using linear mixed-effects models, with metformin concentration, time, and their interaction included as fixed effects, and replicate as a random effect to account for repeated measurements. Pairwise comparisons between treatment groups were corrected for multiple testing using the BH procedure. Different letters indicate statistically significant differences between groups (FDR≤0.05).

**Fig S4:**
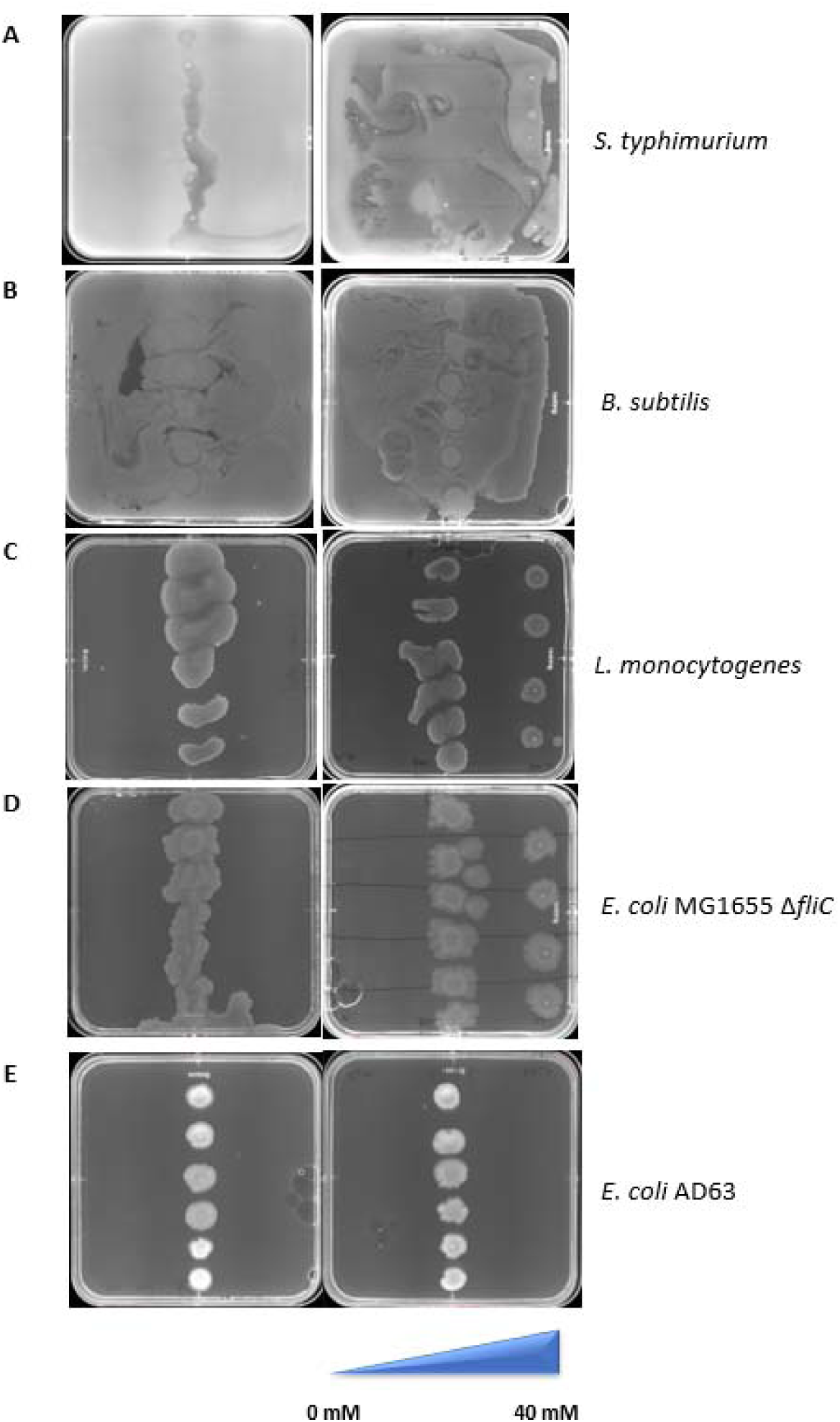
Bacterial migration on soft agar plates in the presence of a metformin gradient. Images of bacterial migration on 0.3% soft agar plates. Plates shown on the left correspond to control conditions without metformin, whereas plates shown on the right contain a metformin gradient ranging from 0 to 40 mM for *S. typhimurium* (A), *B. subtilis* (B), *L. monocytogenes* (C), *E. coli* MG1655 Δ*fliC* (D), *E. coli* AD63 (AD62 deleted in *cheY*) (E). The indicated bacterial strains were inoculated at the center of each plate and incubated at 37°C for all strains except *L. monocytogenes*, incubated at 25°C. Strains were also inoculated in the area of highest concentration of metformin to study the response at high concentrations, except for *B. subtilis*. Differences in migration patterns reflect the bacterial response to the metformin gradient.

**Fig S5:**
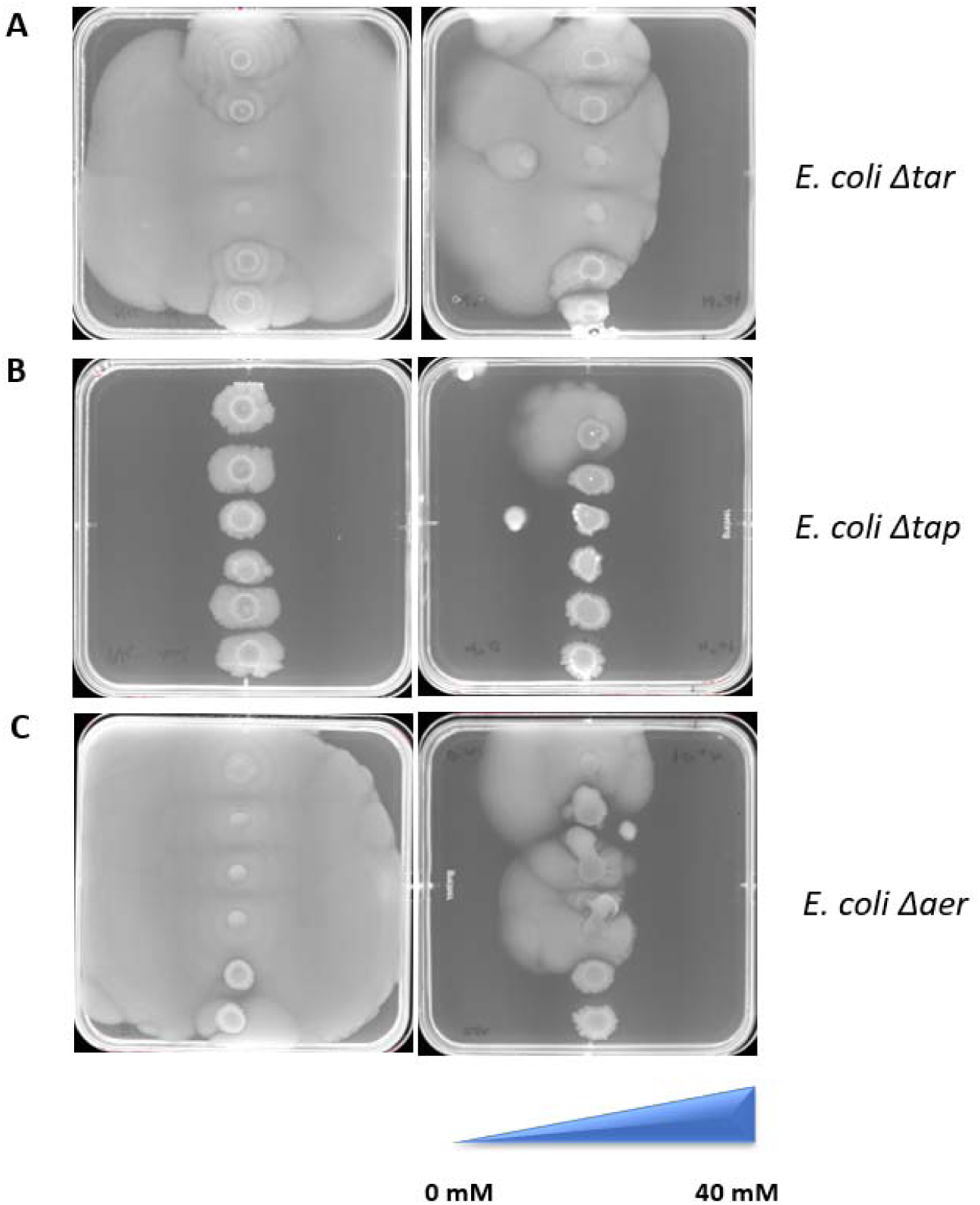
Impact of chemosensor deletion on bacterial migration on soft agar plates in the presence of a metformin gradient. Images of bacterial migration on 0.3% soft agar plates. Plates shown on the left correspond to control conditions without metformin, whereas plates shown on the right contain a metformin gradient ranging from 0 to 40 mM for *E. coli Δtar* (A), *E. coli Δtap* (B) *and, E. coli Δaer* (C). The indicated bacterial strains were inoculated at the center of each plate and incubated at 37°C. Differences in migration patterns reflect the bacterial response to the metformin gradient.

**Fig S6:**
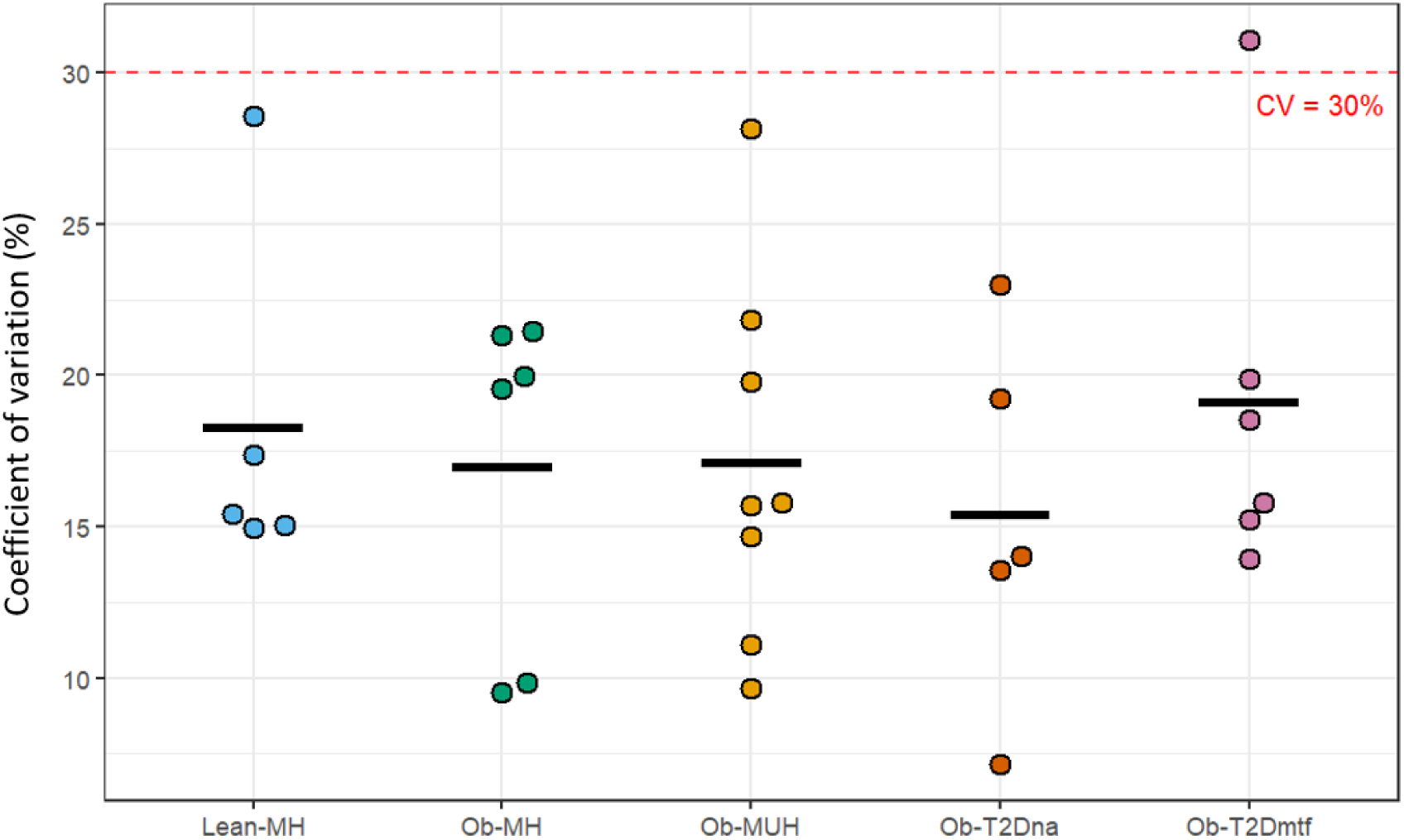
Assessment of analytical reproducibility and biological variability of bacterial penetration with water presented in Fig 3B. Distribution of the coefficients of variation (CV) calculated from experimental replicates for each biological sample. The dashed line indicates the predefined reproducibility threshold (CV = 30%). Technical outliers were removed when necessary to obtain acceptable analytical reproducibility while retaining the maximum number of replicates.

**Table S6:**
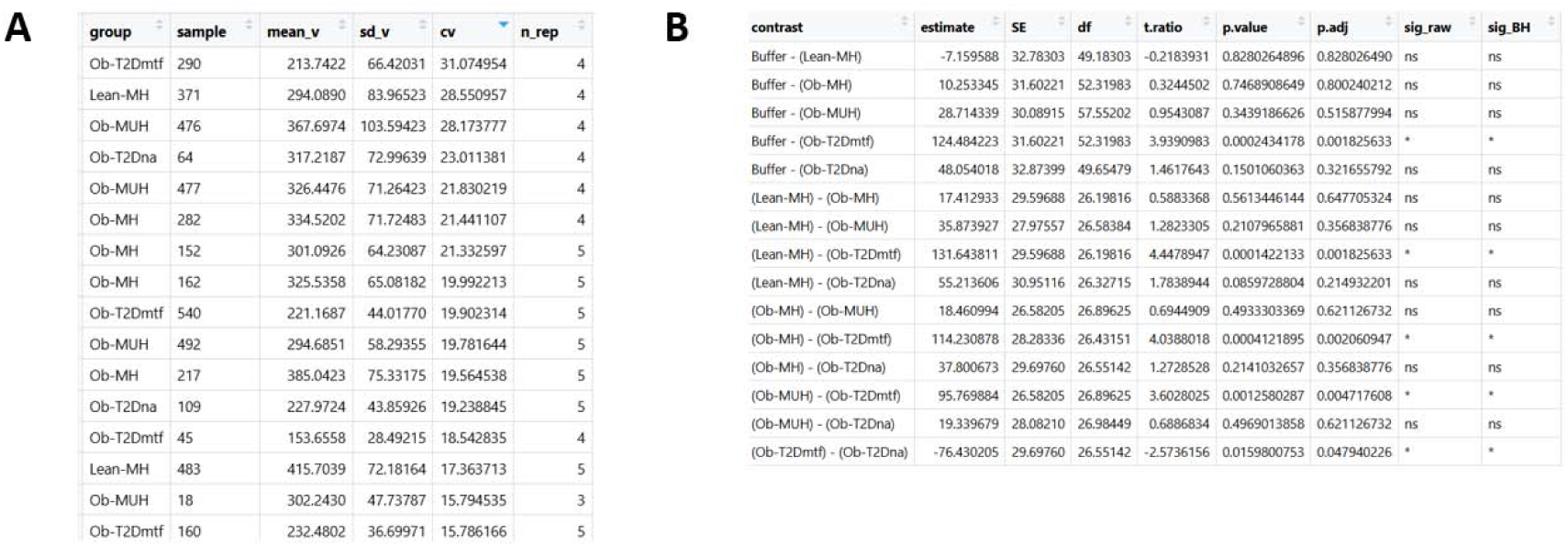
Assessment of analytical reproducibility and biological variability of bacterial penetration with water presented in Fig 3B. **A:** Summary of analytical reproducibility for each experimental group, including the number of biological samples, the number of technical replicates retained after quality control, and the corresponding CV. **B:** Comparison of biological measurements between experimental groups. Following assessment of data normality using the Shapiro–Wilk test, group differences were analyzed using linear mixed-effects models fitted to the raw data, accounting for technical replicates nested within biological samples. Pairwise comparisons were adjusted for multiple testing using the BH procedure.

**Table S7:** Statistical analyses of *E. coli* penetration in mucin in presence of different concentrations of metformin, presented in Fig 3C. Pairwise comparisons of *E. coli* penetration distance in mucin between metformin concentrations at 120 min, derived from one-way ANOVA followed by Tukey’s post-hoc test. Estimate values represent the mean difference in penetration distance between the two compared concentrations, with 95% confidence intervals. Adjusted p-values (Tukey method) are reported for each comparison.

| Group 1 | Group 2 | Estimate | Conf_low | Conf_high | Adjusted p-value |
| --- | --- | --- | --- | --- | --- |
| 0 | 10 | -143.84 | -238.66 | -49.02 | 1.15E-03 |
| 0 | 20 | -190.38 | -285.20 | -95.55 | 2.77E-05 |
| 0 | 30 | -229.18 | -324.01 | -134.36 | 1.43E-06 |
| 0 | 40 | -253.64 | -348.46 | -158.82 | 2.44E-07 |
| 0 | 50 | -313.36 | -408.18 | -218.53 | 4.58E-09 |
| 10 | 20 | -46.54 | -141.36 | 48.29 | 6.57E-01 |
| 10 | 30 | -85.34 | -180.17 | 9.48 | 9.51E-02 |
| 10 | 40 | -109.80 | -204.62 | -14.98 | 1.67E-02 |
| 10 | 50 | -169.51 | -264.34 | -74.69 | 1.45E-04 |
| 20 | 30 | -38.81 | -133.63 | 56.02 | 8.00E-01 |
| 20 | 40 | -63.26 | -158.09 | 31.56 | 3.39E-01 |
| 20 | 50 | -122.98 | -217.80 | -28.16 | 6.05E-03 |
| 30 | 40 | -24.45 | -119.28 | 70.37 | 9.65E-01 |
| 30 | 50 | -84.17 | -178.99 | 10.65 | 1.03E-01 |
| 40 | 50 | -59.72 | -154.54 | 35.11 | 4.00E-01 |

**Supplementary video: Bacterial penetration through mucin depending on the presence and location of metformin** Representative time-lapse of 2 h recording showing the penetration of *E. coli* through a mucin matrix in the absence of metformin (left), with 50 mM of metformin in the mucin (middle) or with 50 mM of metformin in the media (right). Each dot corresponds to a single bacterium.

**Fig S7:**
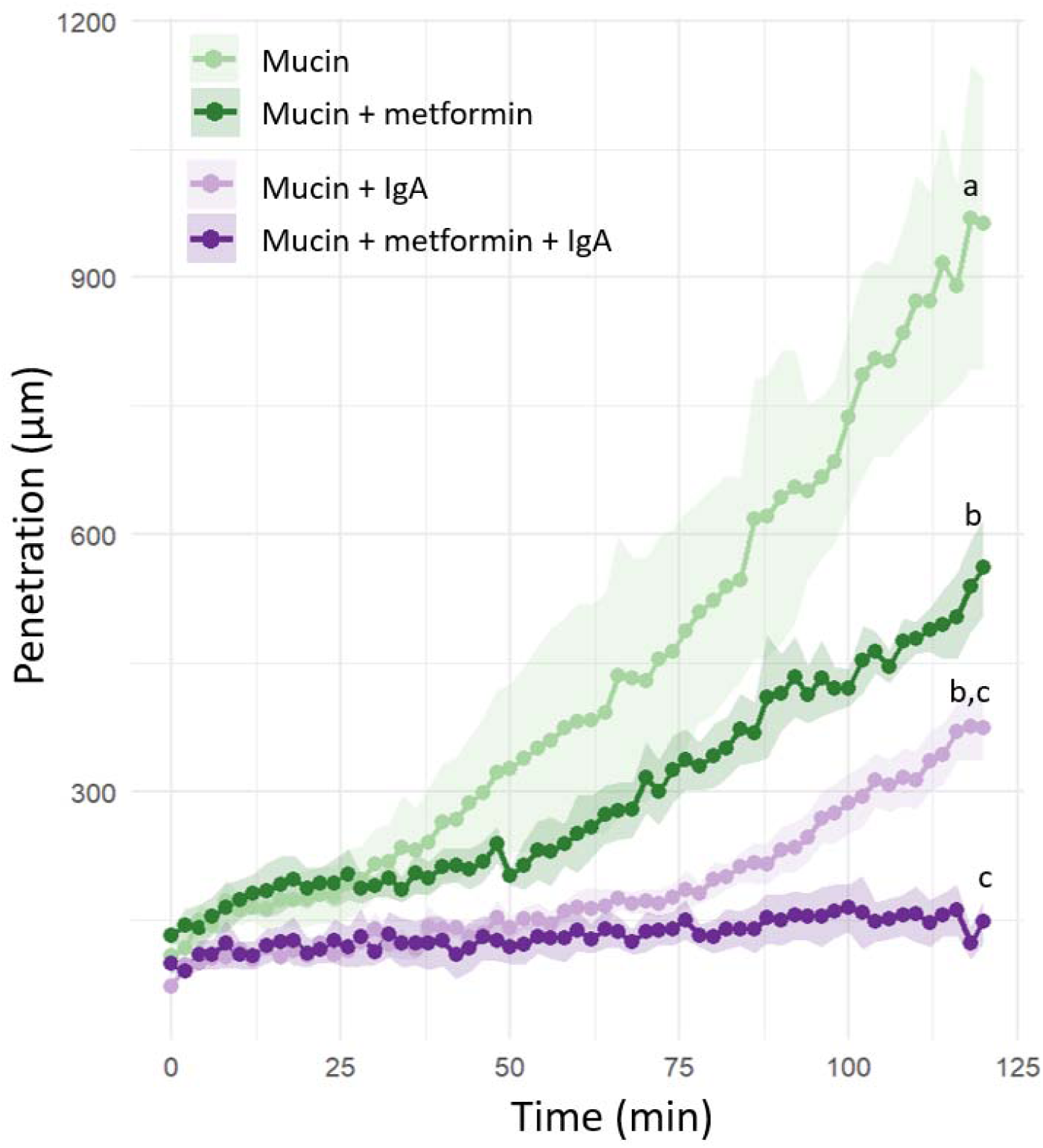
*E. coli* penetration in mucin supplemented with metformin and IgA. Temporal evolution of *E. coli* penetration length in a barrier of mucin without or with metformin (50 mM) and/or IgA (0.077 µg/mL). Curves represent the mean ± SEM of independent biological replicates for each condition (green: without IgA; purple: with IgA; light shades: without metformin; dark shades: with metformin). Statistical significance was assessed using a linear mixed-effects model (Time × Condition, with biological replicate as a random effect), followed by pairwise comparisons between conditions (Tukey-adjusted). Conditions sharing the same letter are not significantly different (pvalue > 0.05); conditions with different letters differ significantly (pvalue ≤ 0.05).

**Table S8:** Statistical analyses of bacterial penetration in mucin in presence of metformin and/or IgA, presented in Fig S7. Statistical significance was assessed using a linear mixed-effects model (Time × Condition, with biological replicate as a random effect), followed by pairwise comparisons between conditions (Tukey-adjusted).

| Groups compared | estimate | t.ratio | p.value | Adjusted p.value |
| --- | --- | --- | --- | --- |
| Mucin_BMB - Mucin_IgA_BMB | 251.68 | 4.17 | 3.13E-03 | 1.33E-02 |
| Mucin_BMB - Mucin_IgA_Met_BMB | 305.02 | 5.05 | 9.87E-04 | 4.30E-03 |
| Mucin_BMB - Mucin_Met_BMB | 142.44 | 2.36 | 4.60E-02 | 1.63E-01 |
| Mucin_IgA_BMB - Mucin_IgA_Met_BMB | 53.34 | 0.88 | 4.03E-01 | 8.14E-01 |
| Mucin_IgA_BMB - Mucin_Met_BMB | -109.24 | -1.81 | 1.08E-01 | 3.36E-01 |
| Mucin_IgA_Met_BMB - Mucin_Met_BMB | -162.58 | -2.69 | 2.74E-02 | 1.03E-01 |

**Table S9:** Statistical analyses of bacterial penetration in pig mucus in presence of metformin, presented in Fig 3E. Pairwise comparisons of estimated marginal means of *E. coli* penetration in pig mucus fed with a low (LFD) or a medium-fat diet (MFD) in presence of 50 mM metformin between the four experimental groups (LFD, MFD, LFD + metformin, MFD + metformin), derived from the linear mixed-effects model. Z-ratios and Tukey-adjusted p-values are reported for each comparison.

| Groups compared | Estimate | Z ratio | Adjusted p-value |
| --- | --- | --- | --- |
| LFD - MFD | -60.78 | -5.794 | <0.0001 |
| LFD - LFD + metformin | -3.43 | -0.327 | 0.988 |
| LFD - MFD + metformin | -5.85 | -0.558 | 0.9445 |
| MFD - LFD + metformin | 57.35 | 5.467 | <0.0001 |
| MFD - MFD + metformin | 54.93 | 5.236 | <0.0001 |
| LFD + metformin - MFD + metformin | -2.42 | -0.231 | 0.9957 |

## Supplemental statistics

### Bacterial growth and flagellin analyses

bioactive flagellin levels across metformin concentrations were compared by one-way ANOVA followed by Tukey-adjusted pairwise comparisons. Bacterial growth curves across metformin concentrations were analyzed using linear mixed-effects models, with metformin concentration, time and their interaction included as fixed effects and biological replicate as a random effect to account for repeated measurements over time. Pairwise comparisons were performed using estimated marginal means, with *P* values adjusted using the BH procedure.

### Regarding Bacterial penetration assays

Bacterial penetration into mucin or mucus was analyzed using linear mixed-effects models where appropriate, with biological replicate included as a random effect to account for technical replicates and/or repeated measurements over time. For experiments using human fecal water, linear mixed-effects models were fitted after CV-based quality control and assessment of model assumptions, and pairwise comparisons were adjusted using the BH procedure. For experiments comparing metformin concentrations at 120 min, differences between conditions were assessed by one-way ANOVA followed by Tukey-adjusted pairwise comparisons. For time-course experiments using mucus from pigs fed a low-fat diet (LFD) or moderate-fat diet (MFD), models included time, diet, metformin treatment and their interactions as fixed effects, with biological replicate included as a random effect. Pairwise comparisons were performed using estimated marginal means with Tukey adjustment. For time-course experiments using mucin with or without metformin and IgA, models included time, experimental condition and their interaction as fixed effects and biological replicate as a random effect. Pairwise comparisons were performed using estimated marginal means with Tukey adjustment.

### Human flagellin and anti-flagellin IgA analyses

Estimated marginal means (EMMs) for fecal anti-flagellin IgA levels and flagellin bioactivity were obtained from linear models including clinical group, age and sex. EMMs therefore represent model-adjusted group means accounting for age and sex. Differences between clinical groups in the joint distribution of anti-flagellin IgA levels and flagellin bioactivity were assessed by multivariate analysis of variance (MANOVA) using Pillai’s trace. Pairwise multivariate comparisons were performed using Hotelling’s T² tests, with P values adjusted using the BH procedure.

### Bioinformatic analyses

Analyses were performed using the R packages ggplot2 (v.3.5.1), Tidyr, v1.3.1, Dplyr v1.1.4, Perm v1.0.0.4, Effsize v0.8.1, Coin v1.4.3, Stringr v1.5.1, Writexl v1.5.1, Ggrepel v0.9.6, Grid v4.4.2, RColorBrewer v1.1.3, Tidyverse v2.0.0, Ggpubr v0.6.0, Emmeans v1.10.6, Lmertest v3.2.1, Lme4 v2.0.1 and Rstatix v0.7.3 R packages were used to analyze, visualize and for the aesthetics of data and figures.

