## Supplementary figures and images for "The chemorepellent effect of metformin limits bacterial penetration into mammalian mucus by altering bacterial motility"

### Supplemental Tables

#### Slide 1
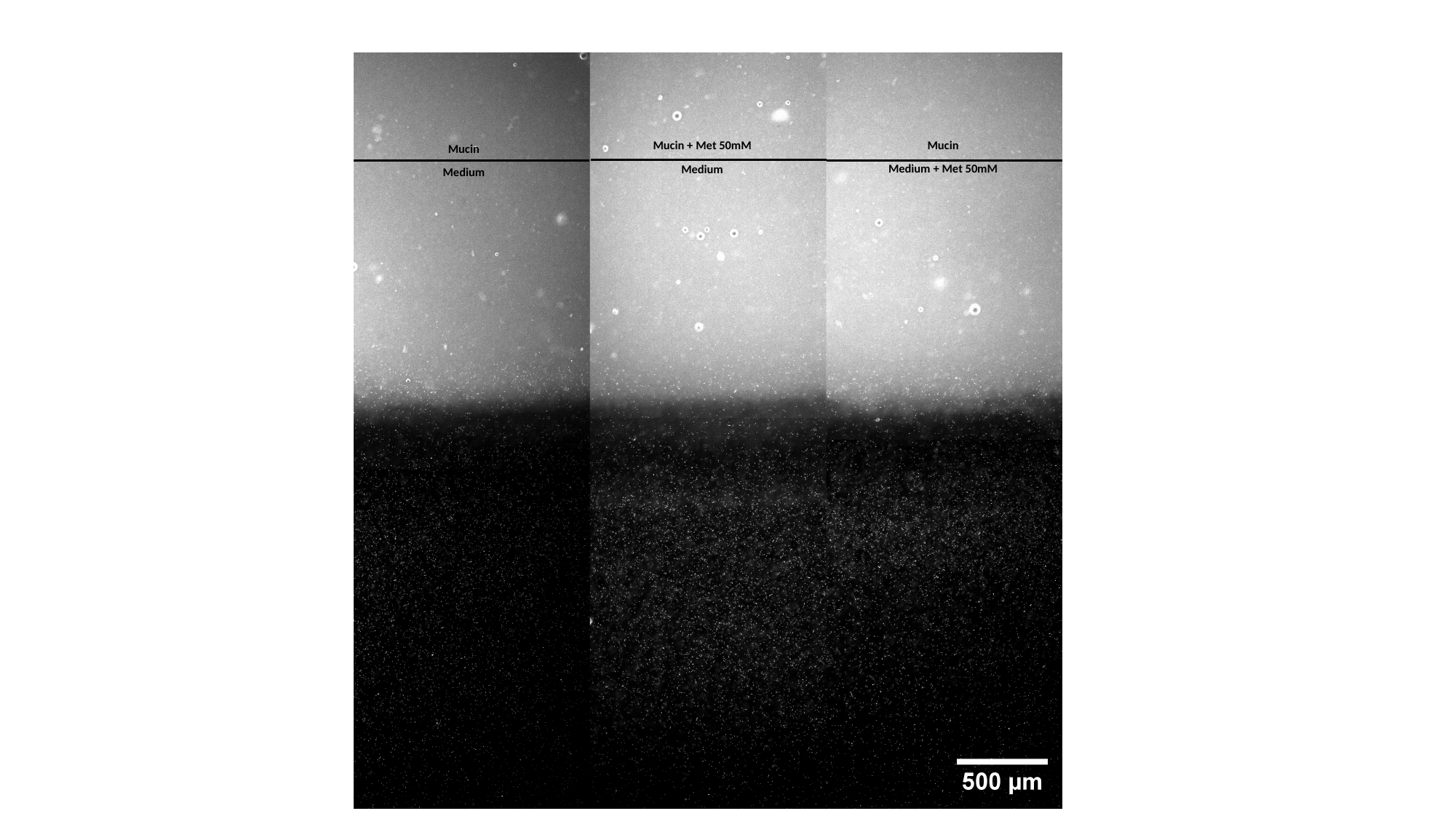

Mucin
Medium + Met 50mM
Mucin + Met 50mM
Medium
Mucin
Medium
